# NFκB and JNK pathways mediate metabolic adaptation upon ESCRT-I deficiency

**DOI:** 10.1101/2024.06.12.598606

**Authors:** Jaroslaw Cendrowski, Marta Wrobel, Michal Mazur, Bartosz Jary, Surui Wang, Michal Korostynski, Anna Dziewulska, Maria Rohm, Agnieszka Dobrzyn, Anja Zeigerer, Marta Miaczynska

## Abstract

Endosomal Sorting Complexes Required for Transport (ESCRTs) are crucial for delivering membrane receptors or intracellular organelles for lysosomal degradation. Yet, how ESCRT dysfunction affects cell metabolism remained elusive. To address this, we analyzed transcriptomes of cells lacking TSG101 or VPS28 proteins, components of ESCRT-I subcomplex. ESCRT-I deficiency reduced the expression of genes encoding enzymes involved in oxidation of fatty acids and amino acids, and increased the expression of genes encoding glycolytic enzymes. Although depletion of ESCRT-I components did not impair mitochondrial biogenesis and ATP-linked respiration it caused intracellular accumulation of lipids and increased lactate production, hallmarks of aerobic glycolysis. Mechanistically, the observed transcriptional reprogramming towards glycolysis in the absence of ESCRT-I occurred due to activation of the canonical NFκB and JNK signaling pathways. Moreover, inhibiting lysosomal activity phenocopied the altered expression of metabolic genes and lipid homeostasis observed for ESCRT-I deficiency, indicating that ESCRT-I restricts glycolysis by mediating lysosomal degradation.

## INTRODUCTION

In order to generate energy, mammalian cells metabolize nutrients, such as glucose, free fatty acids (FFAs) or amino acids, including branched chain amino acids (BCAAs). Typically, these nutrients undergo oxidative metabolism, with initial reactions leading to generation of esters of coenzyme A (CoA), for instance acetylated coenzyme A (acetyl-CoA). These intermediate metabolites are subsequently introduced into the citric acid cycle in mitochondria. Whereas glucose oxidation is often the major source of energy to support cell growth and function [1], in some physiological or pathological conditions cells may prefer to oxidize FFAs or BCAAs [2]. For instance, oxidative break-down of FFAs, known as β-oxidation, is a crucial energy source when glucose supply is limited [3].

Some cell types rely on oxygen-independent phase of glucose metabolism, glycolysis. Glycolysis is the initial chain of reactions of glucose break-down that occurs in the cytoplasm and leads to generation of pyruvate [1]. In cells relying on glycolysis, such as stem cells [4], pyruvate instead of being transformed into acetyl-CoA is converted in the cytosol into lactate, an acidic metabolite that may be released outside the cells to avoid its toxicity [1]. Certain conditions may cause changes in cellular bioenergetics affecting preference for oxidative or glycolytic metabolism, termed metabolic reprogramming. Such reprogramming occurs often in cancer cells (known as Warburg effect) that begin to utilize glycolytic metabolism even in the presence of oxygen (aerobic glycolysis) instead of oxidative metabolism [5, 6]. Another example is inflammation upon which metabolic reprogramming is associated with increased aerobic glycolysis, inhibition of β-oxidation and intracellular accumulation of lipids [7, 8].

Increasing evidence indicates that cellular metabolism strongly depends on the function of lysosomes [9, 10], organelles that degrade different types of cargo delivered from various membrane transport pathways [11]. For instance, lysosomes receive external macromolecules and the plasma membrane proteins via endocytosis (endolysosomal degradation) whereas intracellular cargo including protein aggregates or damaged organelles through macroautophagy (autolysosomal degradation). Endolysosomal degradation of membrane receptors restricts their signaling [12] and this may affect intracellular metabolic cues [13]. Autolysosomal degradation controls the turnover and hence the abundance of metabolic organelles such as mitochondria [14]. Lysosomes are important intracellular source of nutrients including fatty acids or amino acids that come from lysosomal degradation of lipids or proteins [15]. The provision of lysosome-derived nutrients to the cell is sensed by lysosome-associated signaling pathways [16], including signaling of mechanistic Target of Rapamycin Complex 1 (mTORC1) that is activated by lysosome-derived amino acids or cholesterol to promote anabolic and inhibit catabolic pathways [17, 18].

Endosomal sorting complexes required for transport (ESCRTs) may be particularly implicated in regulation of cell metabolism by lysosomes. ESCRTs encompass several protein assemblies (ESCRT-0, I, II, and III) that mediate membrane remodeling in endocytosis, autophagy, cytokinesis, nuclear envelope sealing, and virus budding [19]. Owing to some of their functions, ESCRTs fuel lysosomes with cargo from multiple sources [20], for instance by participating in endolysosomal degradation of many signaling receptors [21], in autolysosomal clearance of protein aggregates [22] and in endolysosomal or autolysosomal degradation of mitochondria [23, 24].

ESCRT dysfunction leads to activation of some stress response pathways (reviewed in [21, 25]). As we previously described, in mammalian cells, inhibition of ESCRT function due to removal of ESCRT-I components, impairs endolysosomal degradation of cytokine receptors [26, 27]. Their endosomal accumulation leads to activation of canonical and non-canonical NFκB signaling, hallmarked by phosphorylation of RELA and accumulation of p52 transcription factors respectively, and induced expression of inflammatory genes [26–28]. Others have shown that ESCRT dysfunction may cause activation of JNK signaling [29, 30]. Canonical NFκB signaling was found to inhibit fatty acid oxidation during cardiac hypertrophy [31] and to promote aerobic glycolysis in sarcoma cells [32], whereas JNK signaling was reported to either inhibit or promote glycolytic metabolism [33]. However, whether the regulation of NFκB or JNK pathways upon ESCRT dysfunction has consequences for cell metabolism has not been addressed.

Our recent studies began to unravel particular metabolic pathways regulated by ESCRT. VPS37A, one of ESCRT-I components was shown to have a specific function in the liver in mediating glucagon receptor degradation, thereby regulating hepatic glucose production [34]. In another study we discovered that by fueling lysosomes with macromolecules from multiple sources, ESCRT-I promotes substrate-specific mTORC1 signaling that inhibits TFEB/TFE3 transcription factors. While ESCRT-I depletion does not affect the general mTORC1 signaling, it leads to activation of TFEB/TFE3 transcription factors that stimulate the expression of genes involved in lysosome biogenesis [11]. Although TFEB/TFE3 signaling may control lipid metabolism and mitochondria biogenesis [35, 36], it remains unknown whether activation of these transcription factors affects cell metabolism upon ESCRT deficiency.

Here, we aimed at studying metabolic consequences of ESCRT-I deficiency. By a transcriptomic approach and its subsequent validation, we discovered that ESCRT-I depletion causes a transcriptional reprogramming towards glycolytic metabolism that occurs due to activation of NFκB and JNK signaling pathways and likely as a result of inhibited lysosomal degradation.

## RESULTS

### Lack of ESCRT-I leads to reduced expression of genes involved in oxidative metabolism of carboxylic acid-containing molecules such as amino acids and fatty acids

We have previously shown that in HEK293 cells, lack of ESCRT-I causes a very prominent activation of inflammatory NFκB-dependent signaling [27] and lysosomal starvation-related TFEB/TFE3-dependent signaling [11]. However, whether removing ESCRT-I alters the expression of metabolic genes in these cells has not been investigated. Hence, we performed a microarray analysis of HEK293 cells in which we depleted the ESCRT-I components, TSG101 or VPS28 proteins using two distinct siRNAs (designated #1 and #2) against each component (**Fig. 1A**) and analyzed the cells three days post transfection (3 dpt). Depletion of one of these proteins led to removal of the other one (**Fig. 1A**), consistent with destabilization of the whole complex that occurs upon depletion of any of its core components [28, 37].

**Figure 1.**
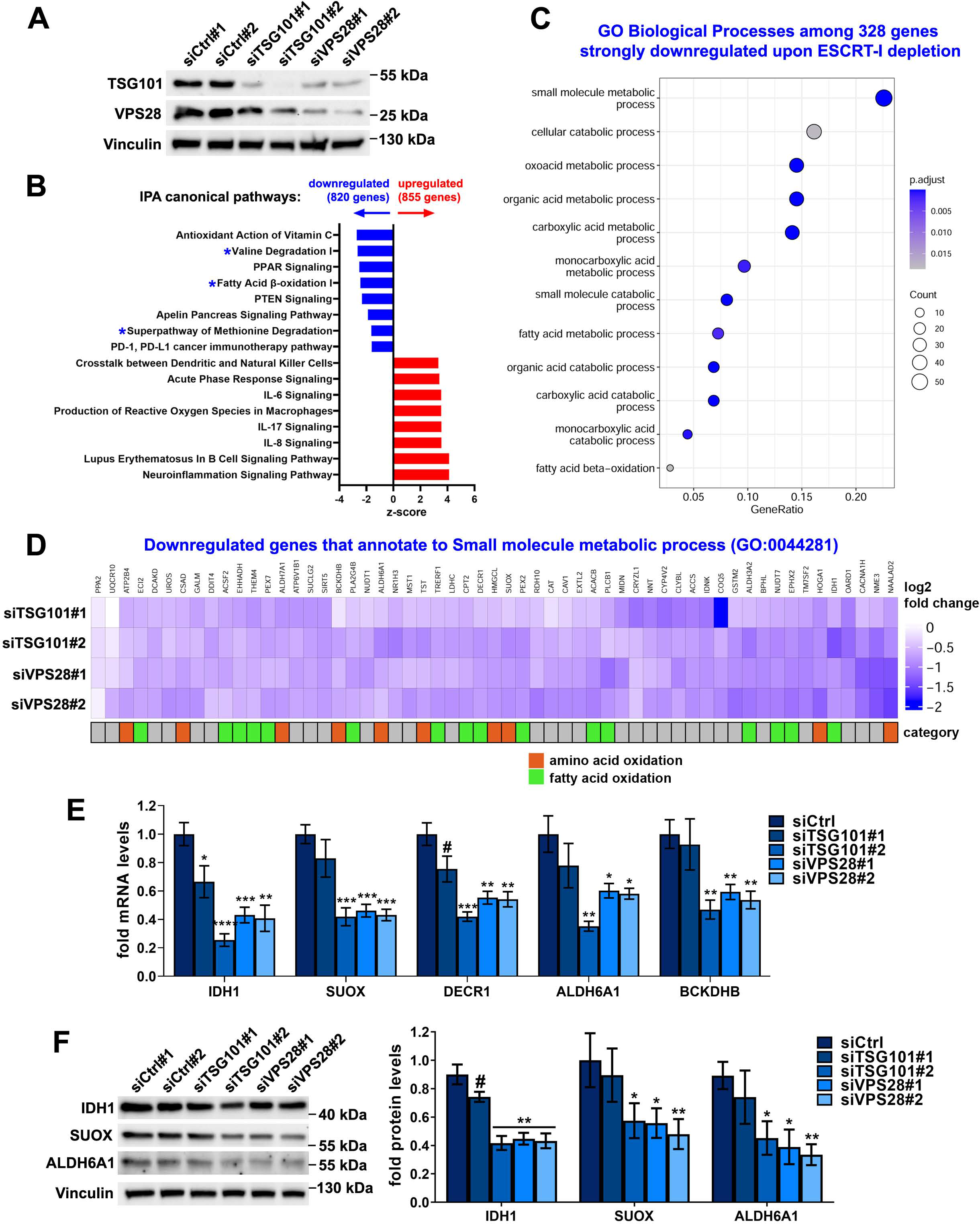
ESCRT-I dysfunction leads to reduced expression of genes involved in oxidative metabolism of amino acids and fatty acids. (A) Western blots showing the depletion efficiencies of ESCRT-I subunits, TSG101 or VPS28 using two single siRNAs for each component (siTSG101#1 or siTSG101#2, siVPS28#1 or siV2PS8#2), as compared to control cells (treated with non-targeting siRNAs, Ctrl#1 or #2) in HEK293 cells. Vinculin used as a gel loading control. **(B)** Ingenuity Pathway Analysis (IPA) of microarray results, showing top canonical pathways identified by annotation of genes whose expression was significantly (FDR < 0.05) downregulated or upregulated in HEK293 cells depleted of ESCRT-I using two single siRNAs for each component, as compared to control cells. Blue asterisks indicate annotations related to metabolism of amino acids and fatty acids. Microarray data analysis was performed based on three independent experiments. **(C)** Gene ontology (GO) analysis of top biological processes identified by annotation of genes detected in microarray experiments as those with strongly downregulated expression (log2 fold change ≤ -0.6; FDR < 0.05) upon ESCRT-I removal. **(D)** Heatmap visualizing expression of genes annotated to “small molecule metabolic process” (GO:0044281), whose mRNA levels were detected by microarray as downregulated after ESCRT-I removal. The genes encoding enzymes involved in oxidation of amino acids or fatty acids indicated with orange and green rectangles. **(E)** qPCR results showing the expression of genes encoding the indicated oxidative metabolism enzymes in HEK293 cells lacking ESCRT-I, as compared to control cells, presented as fold changes with respect to averaged values measured for siCtrl#1 and #2 cells (siCtrl). Mean values (n = 5 ± SEM) are presented. **(F)** Representative western blots (left panel) showing the levels of the indicated oxidative enzymes in control or ESCRT-I-deficient cells. The graph (right panel) shows protein levels as fold change with respect to averaged values measured for control cells (siCtrl) by densitometry analysis of western blotting bands. Vinculin was used as a gel loading control. Values derived from independent experiments and their means (n = 4 ± SEM) are presented. All the analyses shown in A-F were performed at three days post transfection with siRNAs (3 dpt). Statistical significance tested by comparison to siCtrl. ^#^P < 0.1, *P < 0.05, **P < 0.01, ***P < 0.001, ****P < 0.0001.

Among genes whose expression was commonly altered in samples with TSG101 or VPS28 depletion, we detected 855 genes with significantly upregulated expression and 820 with significantly downregulated expression, as compared to control cells (**Fig. 1B**). These genes were functionally annotated using the Ingenuity Pathway Analysis. As expected from our previous studies [11, 26, 27], ESCRT-I depletion led to elevated expression of many genes involved in inflammation and stress response (**Fig. 1B**).

Importantly, we observed that among genes with reduced expression in cells lacking ESCRT-I were those annotated to oxidative breakdown of amino acids and fatty acids (**Fig. 1B**). Our further analysis using Gene Ontology Biological Processes database, focusing only on 328 genes with the strongest downregulation (log2 fold change ≤ -0,6), showed that ESCRT-I depletion led to reduced expression of over 50 genes involved in “small molecule metabolism” (**Fig. 1C-D**). This list included mainly genes encoding enzymes responsible for oxidative metabolism of carboxylic acid-containing molecules such as fatty acids and amino acids (**Fig. 1D, Table S1**). Given that these genes encode enzymes involved in oxidation of such molecules, we refer to them hereafter as “amino acid or fatty acid oxidation genes”.

To validate the transcriptomic results, we focused on genes encoding IDH1 and DECR1 enzymes, that participate in fatty acid metabolism [38, 39], as well as SUOX, ALDH6A1 and BCKDHB [40–42] that are involved in amino acid metabolism. By quantitative RT-PCR (qRT-PCR), we independently confirmed the reduced mRNA levels of each of these enzymes in HEK293 cells lacking TSG101 or VPS28 (**Fig. 1E**). However, siTSG101#1, that was less efficient in depleting TSG101 and VPS28 proteins than siTSG101#2 (shown in **Fig. 1A**) had a weaker effect on the mRNA levels of the analyzed enzymes. To investigate whether reduced expression of the chosen genes occurred also in other cell types, we performed qRT-PCR analysis in HepG2 hepatoblastoma cell line with siRNA-mediated removal of TSG101 or VPS28 proteins using siTSG101#2 and siVPS28#1 (**Fig. S1A**) or CRISPR-Cas9-mediated knock-out of *TSG101* gene (**Fig. S1B-C**). The efficiencies of using siRNAs or CRISPR-Cas9 system in these cells were assessed by qRT-PCR (**Fig. S1A**) or western blotting (**Fig. S1B**), respectively. In each case, deficiency of ESCRT-I complex led to a reduced expression of analyzed amino acid or fatty acid oxidation genes (**Fig. S1A and C**).

To address whether changes in the expression of amino acid or fatty acid oxidation genes observed upon ESCRT-I deficiency could affect cell metabolism, we tested protein levels of chosen enzymes in HEK293 cells by western blotting. We confirmed that siRNA-mediated depletion of TSG101, using siTSG101#1, or VPS28, using siVPS28#1 or #2, at 3 dpt caused reduced abundance of IDH1, SUOX and ALDH6A1 oxidative enzymes (**Fig. 1F**). Thus, the transcriptomic analysis and its validation demonstrated that ESCRT-I deficiency causes reduced gene expression and protein levels of enzymes involved in oxidation of small molecules such as amino acids and fatty acids.

### ESCRT-I deficiency causes intracellular accumulation of lipids

Reduced expression of genes encoding enzymes of fatty acid oxidation is often associated with intracellular accumulation of lipids [43]. Moreover, we observed increased expression of several genes encoding enzymes of biosynthesis of various types of lipids (fatty acids, triglycerides or phospholipids) in cells lacking ESCRT-I (**Fig. S2**). Thus, we investigated the abundance of such lipids in cells with TSG101 or VPS28 depletion using single siRNAs (siTSG101#2 and siVPS28#2) against each protein (**Fig. 2A**).

**Figure 2.**
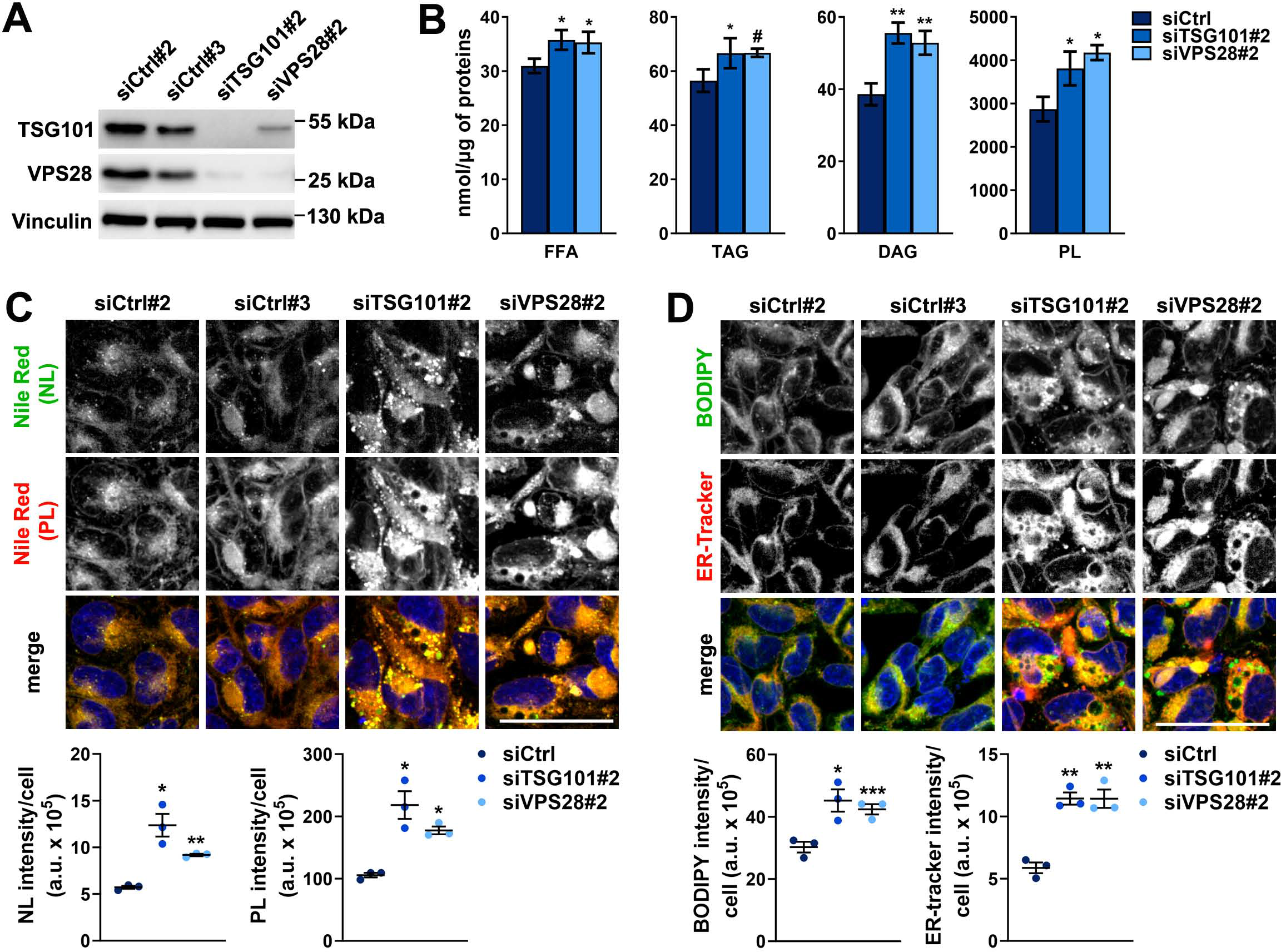
Cells lacking ESCRT-I have elevated levels of free fatty acids (FFAs) and fatty acid- containing lipids. (A) Western blots showing the efficiencies of siRNA-mediated depletions of ESCRT-I subunits, TSG101 or VPS28 (cells treated with siTSG101#2 or siV2PS8#2), as compared to control conditions (two non-targeting siRNAs, Ctrl#2 or #3) in HEK293 cells. Vinculin used as a gel loading control. **(B)** Results of gas chromatography followed by mass spectrometry (GC-MS) showing Intracellular levels of free fatty acids - FFA, triglycerides - TG, diacylglycerides - DAG or phospholipids - PL (shown as nmol/µg of proteins) in control or ESCRT-I-depleted cells. Values derived from independent experiments and their means (n = 4 ± SEM) are presented. Values for siCtrl are averaged values measured for cells transfected with siCtrl#2 or siCtrl#3. **(C-D)** Maximum intensity projection confocal images of live control or ESCRT-I-depleted cells. The images show the intracellular distribution of neutral lipids – NL (green), or phospholipids – PL (red) stained with Nile Red dye (shown in C) as well as the intracellular distribution of NLs stained with BODIPY 493/503 and the ER stained with ER-tracker (green or red, respectively in D). Cell nuclei marked with Hoechst stain (blue). Scale bar, 50 μm. The dot plots on the bottom show total fluorescence intensities per cell (expressed in arbitrary units, a.u.), as compared to averaged values measured for cells transfected with siCtrl#2 or siCtrl#3 (siCtrl). Values derived from independent experiments (dots) and their means (n = 3 ± SEM) are presented. All the analyses shown in B-D were performed at three days post transfection with siRNAs (3 dpt). Statistical significance tested by comparison to the siCtrl values. ^#^P < 0.1, *P < 0.05, **P < 0.01, ***P < 0.001.

To assess the intracellular content of fatty acids, we performed gas chromatography followed by mass spectrometry (GC-MS) (**Fig. 2B**). We measured free fatty acids (FFAs) and fatty acid-containing lipids, i.e., triglycerides (TGs), diacylglycerides (DAGs) and phospholipids (PLs). Depletion of TSG101 or VPS28 led to slightly elevated levels of FFAs and TGs and to even stronger increase in membrane lipids, DAGs and PLs (**Fig. 2B**). To verify elevated lipid levels, we stained control and ESCRT-I-depleted cells with Nile Red dye (NR) and imaged them by confocal microscopy. NR emits green fluorescence, when bound to neutral lipids (NLs, that include cholesterol or TGs), or red fluorescence, when bound to PLs [44]. Consistent with the GC-MS results, we observed that lack of ESCRT-I led to increased levels of both NLs and PLs (**Fig. 2C**). We confirmed elevated abundance of NLs in cells lacking TSG101 or VPS28 by using BODIPY 493/503 dye (**Fig. 2D**) that emits green fluorescence when bound to these lipids [45]. The amounts of NLs were increased primarily in lipid droplets – LDs, identified as punctate structures positive for both NLs and PLs [46], that were more abundant and larger upon ESCRT-I depletion (**Fig. 2C-D**). PLs accumulated in ESCRT-I-deficient cells also outside of LDs, in a region resembling the ER by shape and localization (**Fig. 2C**). As PLs are the most abundant lipids in the cell, that build cellular membranes, we reasoned that their accumulation in the ER region could reflect increase of ER volume. Indeed, we noticed that cells lacking TSG101 or VPS28 had a strongly increased ER content as measured by confocal microscopy using ER-tracker Red dye (**Fig. 2D**), although the functional significance of this observation is unclear.

Hence, we discovered that lack of TSG101 or VPS28 leads to increased abundance of FFAs and lipids, such as PLs that accumulate in intracellular membranes including the enlarged ER. Although increased amount of fatty acid-containing lipids in cells lacking ESCRT-I could be in part due to their impaired lysosomal degradation, it could also result from reduced oxidative breakdown of FFAs and increased lipid biosynthesis, as implied by the changes in gene expression described above (**Fig. 1** **and S2**).

### ESCRT-I deficiency does not impair mitochondrial biogenesis or ATP-linked mitochondrial respiration

By GO Cellular Component analysis of genes with strongly reduced expression in cells lacking TSG101 or VPS28, we identified many genes that encode mitochondrial proteins (**Fig. S3**, **Table S2**; around 50 genes that included many of the amino acid or fatty acid oxidation genes indicated in Fig. 1D). Reduced expression of genes encoding mitochondrial proteins could potentially affect mitochondrial biogenesis. However, we did not observe a general reduction in the expression of genes encoding structural mitochondrial proteins or core components of the citric acid cycle or oxidative phosphorylation. To investigate the consequence of ESCRT-I deficiency on the abundance of mitochondria, we performed confocal microscopy analysis of HEK293 cells stained with MitoTracker Deep Red FM dye. We did not observe reduced mitochondrial content in the absence of ESCRT-I at 3 dpt (**Fig. 3A**). Conversely, we observed an increase in mitochondrial abundance, particularly strong in cells lacking TSG101 (**Fig. 3A**). These data indicated that ESCRT-I deficiency did not impair the biogenesis of mitochondria.

**Figure 3.**
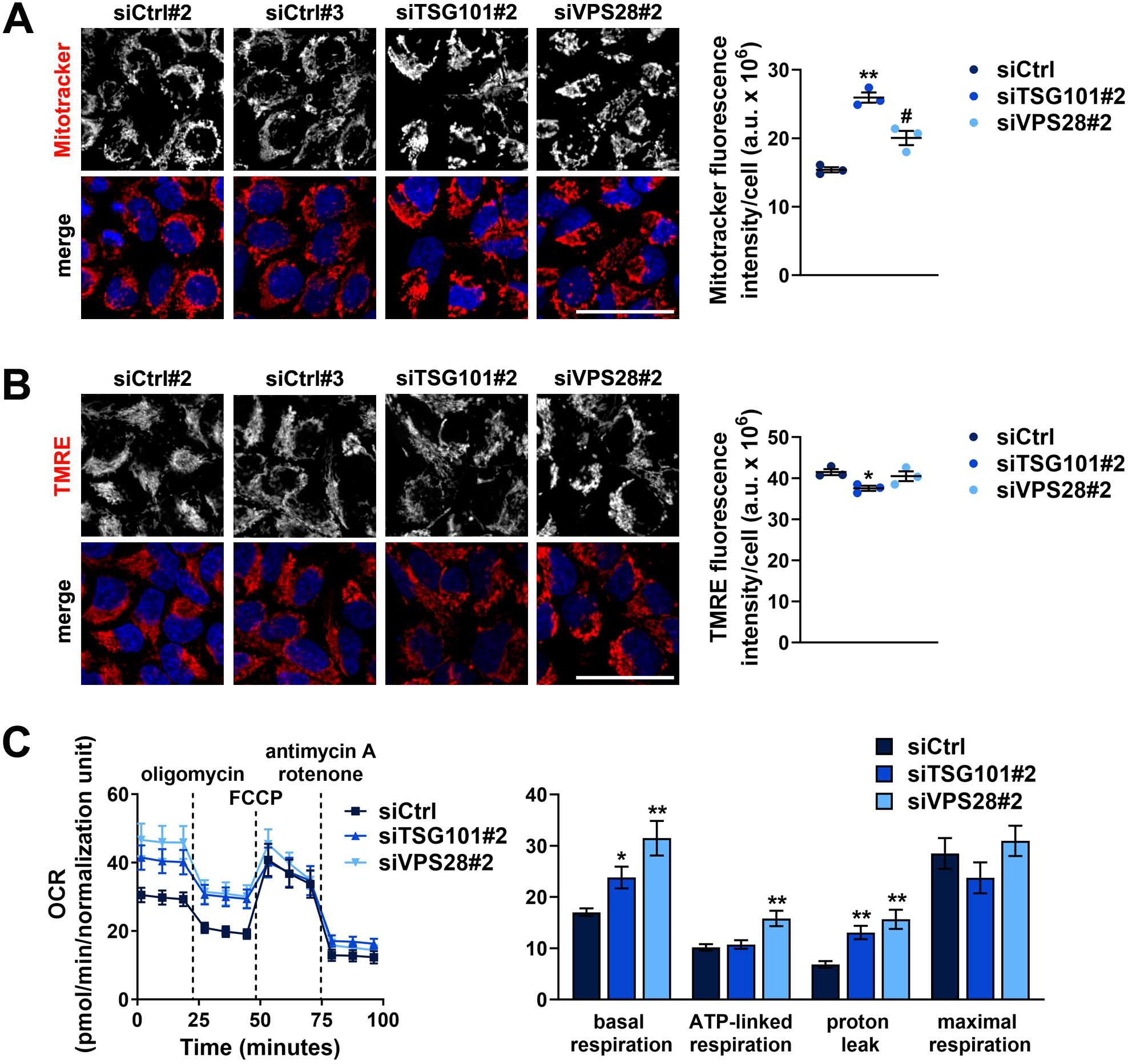
The abundance of functional mitochondria and the rate of ATP-linked mitochondrial respiration are not reduced upon ESCRT-I deficiency. (A-B) Maximum intensity projection confocal images of live control (treated with non-targeting siRNAs, Ctrl#2 or #3) or ESCRT-I-depleted cells (treated with siTSG101#2 or siV2PS8#2). The images show the intracellular content and distribution of all mitochondria stained with MitoTracker Deep Red FM dye (red colour in A) and of mitochondria with proper membrane potential stained with Tetramethylrhodamine, Ethyl Ester (TMRE, red, shown in B). Cell nuclei marked with Hoechst stain (blue). Scale bar, 50 μm. The dot plots show total fluorescence intensities per cell (expressed in arbitrary units, a.u.), as compared to averaged values measured for cells transfected with siCtrl#2 or siCtrl#3 (siCtrl). Values derived from independent experiments (dots) and their means (n = 3 ± SEM) are presented. **(C)** A representative time-series graph (left) showing changes of oxygen consumption rate (OCR) with time (pmol/min) in control or ESCRT-I-depleted cells upon: basal respiration (untreated cells; time-points 1-3), inhibition of ATP- linked respiration (oligomycin treatment; time-points 4-6), maximal respiration (FCCP treatment; time-points 7-9) and non-mitochondrial respiration (antimycin A and rotenone treatment; time- points 10-12). The bar graph (right) shows the intensity of indicated processes calculated based on the results shown in the time-series graph. The results shown in both graphs were normalized to cell number reflected by DNA staining with Hoechst 33342 dye. Mean values in both graphs derived from technical repetitions of one experiment (n = 4 or 5 ± SEM) are presented. All the analyses shown in A- C were performed at three days post transfection with siRNAs (3 dpt). Statistical significance tested by comparison to the siCtrl values. ^#^P < 0.1, *P < 0.05, **P < 0.01.

As ESCRTs are required for mitochondria degradation in lysosomes [23, 24], the accumulation of mitochondria observed in cells lacking TSG101 or VPS28 could be a result of impaired autophagic removal of damaged mitochondria. Hence, to address whether ESCRT-I deficiency affects the functionality of mitochondria, we measured mitochondrial membrane potential using Tetramethylrhodamine Ethyl Ester (TMRE) dye that stains properly functioning mitochondria. By confocal microscopy, we observed that the TMRE fluorescence intensity was slightly reduced upon TSG101 depletion and not altered upon VPS28 depletion (**Fig. 3B**). Hence, the overall mitochondrial membrane potential did not increase upon ESCRT-I deficiency (**Fig. 3B**) as it was the case for overall mitochondria content (shown in **Fig. 3A**). This analysis indicated that cells lacking ESCRT-I retain similar amounts of functional mitochondria as in control cells but accumulate excess mitochondria that do not have proper membrane potential.

To address whether the reduced expression of amino acid or fatty acid oxidation genes or the accumulation of damaged mitochondria upon ESCRT-I deficiency affect mitochondrial respiration, we measured oxygen consumption rate (OCR) using the Agilent Seahorse XFe24 Analyzer. Control HEK293 cells showed typical OCR curve, indicating basal respiration (as a sum of ATP-linked and proton leak-related processes) that was lower than maximal respiratory capacity (**Fig. 3C**). However, cells lacking ESCRT-I components had elevated basal respiration, reaching maximal capacity, due to increased proton leak (**Fig. 3C**). The elevated proton leak was consistent with the above described accumulation of damaged mitochondria upon ESCRT-I deficiency (shown in Fig. 3A-B). Importantly, the OCR linked to ATP production was not impaired in cells lacking ESCRT-I depletion (**Fig. 3C**). It was not affected by depletion of TSG101 and was even elevated upon VPS28 depletion as compared to control cells (**Fig. 3C**).

Overall, although lack of TSG101 or VPS28 leads to accumulation of damaged mitochondria it does not affect the abundance of mitochondria with proper membrane potential. Intriguingly, despite strongly impaired lysosomal degradation of proteins [11, 27] and lipids (shown in Fig. 2B-C), as well as reduced levels of enzymes involved in amino acid and fatty acid degradation (shown in Fig. 1F), ESCRT-I depletion does not impair ATP-dependent mitochondrial respiration. These results suggested that cells lacking ESCRT-I may reprogram their metabolism from oxidizing lysosome-derived nutrients towards utilizing other energy sources.

### Cells lacking ESCRT-I activate glycolytic metabolism

FFAs or BCAAs may serve as alternative energy sources to glucose metabolism [8]. Hence, we were intrigued whether the reduced expression of amino acid or fatty acid oxidation genes in ESCRT-I-deficient cells was associated with changes in expression of genes involved in glucose catabolism. Although the Ingenuity Pathway Analysis of genes with commonly induced expression upon TSG101 or VPS28 depletion (**Fig. 1B**) did not indicate such annotation, we investigated in more detail the transcriptomic results focusing on genes encoding enzymes involved in glycolysis, hence metabolism of glucose to pyruvate (**Fig. 4A**), and in conversion of pyruvate to acetyl-CoA (**Fig. 4B**). We found that ESCRT-I deficiency caused elevated expression of several genes that encode glycolytic enzymes (**Fig. 4A**) but had no particular effect on the expression of genes responsible for pyruvate to acetyl-CoA conversion (**Fig. 4B**). By qRT-PCR analysis performed at 3 dpt (the same time-point as of the transcriptomic analysis), we verified the increased expression of HK2, ENO2, PFKFB3 and PFKFP glycolytic enzymes in HEK293 cells upon depletion of VPS28 with two independent siRNAs (**Fig. 4C**). However, such increase was not observed in cells transfected with siTSG101#1 and occurred only for ENO2 and PFKFB3 in cells transfected with siTSG101#2 (**Fig. 4C**).

**Figure 4.**
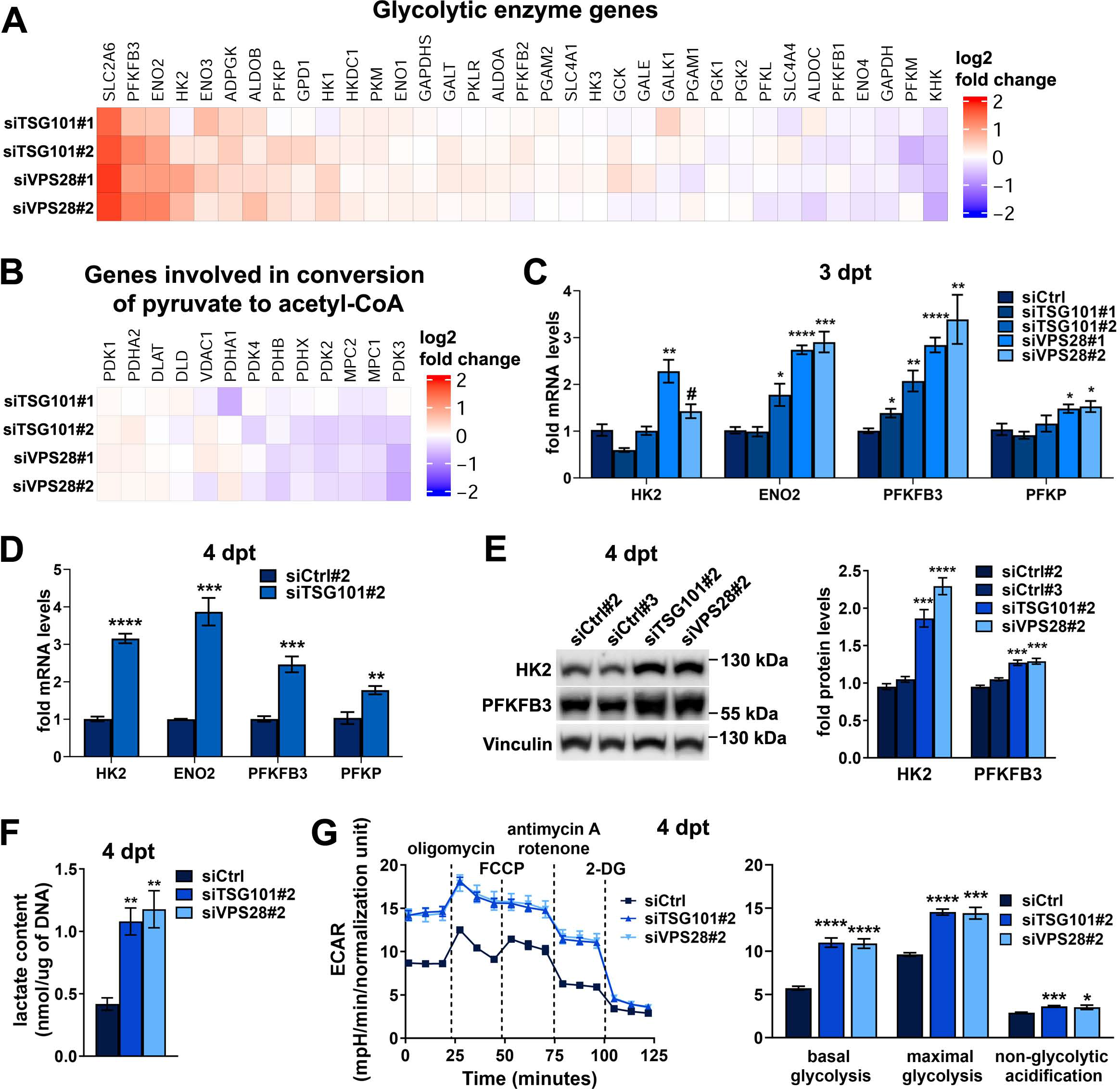
ESCRT-I dysfunction leads to increased glycolysis. (A-B) Heatmaps visualizing microarray results regarding the expression of genes encoding enzymes involved in glycolysis (in A) or in production of acetyl-CoA from pyruvate (in B) in HEK293 cells after removal of ESCRT-I using two siRNAs for each component (siTSG101#1 or siTSG101#2, siVPS28#1 or siVPS28#2), as compared to control cells (treated with non-targeting siRNAs, Ctrl#1 or #2). Microarray data analysis was performed based on three independent experiments at three days post transfection with siRNAs (3 dpt). **(C)** qPCR results showing the expression of genes encoding the indicated glycolytic enzymes at 3 dpt in cells with ESCRT-I removal, as compared to control cells. The mean values (n = 4 ± SEM) are presented as fold changes with respect to average values for control cells (siCtrl). **(D)** qPCR results showing the expression of genes encoding glycolytic enzymes in HEK293 cells depleted of TSG101 using a single siRNA (siTSG101#2), as compared to control cells (treated with non-targeting siRNA, Ctrl#2). The mean values (n = 4 ± SEM) presented as fold changes with respect to values for control cells at 4 dpt. **(E)** Representative western blots (left panel) showing the levels of the indicated glycolytic enzymes in control cells (treated with non-targeting siRNAs, Ctrl#2 or #3) or ESCRT-I- depleted (siTSG101#2 or siVPS28#2). The graph (right panel) shows protein levels as fold change with respect to averaged values measured for siCtrl#2 and #3 assessed by densitometry analysis of western blotting bands. Vinculin was used as a gel loading control. Mean values (n = 4 ± SEM) are presented. **(F)** Intracellular levels of lactate (shown as nmol/µg of DNA) in control (siCtrl#1 or #2) or ESCRT-I-deficient cells. Mean values (n = 3 ± SEM) are presented. Values for siCtrl are averaged values measured for cells transfected with siCtrl#1 or siCtrl#2. **(G)** A representative time-series graph (left) showing changes of extracellular acidification rate (ECAR) with time (mpH/min) in control (siCtrl#2 or #3) or ESCRT-I-depleted cells at 4 dpt upon: basal respiration (untreated cells; time-points 1-3), nhibition of ATP-linked respiration (oligomycin treatment; time-points 4-6), maximal respiration (FCCP treatment; time-points 7-9), non-mitochondrial respiration (antimycin A and rotenone treatment; time-points 10-12) and inhibition of glycolysis (2-DG treatment; time-points 13-15). The bar graph (right) shows the intensity of indicated processes calculated based on the results shown in the time-series graph. The results shown in both graphs were normalized to cell number reflected by DNA staining with Hoechst 33342 dye. Mean values in both graphs derived from technical repetitions of one experiment (n = 5 ± SEM) are presented. Statistical significance in C and F tested by comparison to siCtrl, whereas in D and E tested by comparison to siCtrl#2. ^#^P < 0.1, *P < 0.05, **P < 0.01, ***P < 0.001, ****P < 0.0001.

Reasoning that 3 dpt could be too early to observe a full effect on glycolytic gene expression in cells lacking TSG101, we performed the analysis in cells transfected with siTSG101#2 at 4 dpt and observed a prominent increase in the expression of all analyzed glycolytic genes (**Fig. 4D**). The upregulated expression of genes encoding ENO2, PFKFB3 and PFKP (but not HK2) also occurred in ESCRT-I-deficient HepG2 cells at 3 dpt (**Fig. S4A-B**). As in HEK293 cells, we observed a stronger increase in the expression of these glycolytic genes in HepG2 cells with siRNA-mediated VPS28 depletion than upon TSG101 depletion (**Fig. S4A**). However, CRISPR-Cas9-mediated TSG101 depletion led to clearly upregulated expression of genes encoding ENO2, PFKFB3 and PFKP (**Fig. S4B**).

Next, we addressed whether increased expression of genes encoding particular glycolytic enzymes translates into their higher abundance and increased glycolytic metabolism. Analyzing the cells at 4 dpt, we observed elevated protein levels of HK2 and PFKFB3 enzymes upon depletion of both VPS28 or TSG101 (**Fig. 4E**). To address whether ESCRT-I-deficient HEK293 cells activate glycolytic metabolism, we measured the intracellular content of lactate, the product of anaerobic glucose metabolism. We found that at 4 dpt, cells lacking TSG101 or VPS28 had clearly elevated intracellular levels of lactate (**Fig. 4F**). To verify the effect of ESCRT-I deficiency at 4 dpt on glycolysis, we used the Agilent Seahorse XFe24 Analyzer to measure the extracellular acidification rate (ECAR) that is increased upon release of lactate from cells that undergo glycolysis [47]. This analysis showed that, as compared to control cells, cells lacking TSG101 or VPS28 had elevated ECAR, primarily due to glycolysis (**Fig. 4G**), although their non-glycolytic acidification was also slightly elevated (**Fig. 4G**). Of note depletion of ESCRT-I components caused an increase of both, basal glycolysis as well as maximal glycolytic capacity (**Fig. 4G**).

Hence, in cells lacking ESCRT-I, the inhibited expression of amino acid or fatty acid oxidation genes is associated with activated expression of glycolytic genes and induction of glycolytic metabolism. These results suggest that functional ESCRT-I may promote oxidative metabolism of lysosome-derived nutrients over glycolytic metabolism.

### mTORC1 signaling is not implicated in regulation of cell metabolism upon ESCRT-I depletion

Next, we sought to investigate which signaling pathways could be involved in the altered expression of metabolic genes upon ESCRT-I depletion. Cell metabolism is largely controlled by mTORC1 signaling that regulates amino acid, fatty acid and glucose metabolism. We have previously found that in RKO colorectal cancer cells, ESCRT-I deficiency does not affect general mTORC1 signaling, for instance phosphorylation of mTOR kinase target S6K1 [11]. To verify whether general mTORC1 signaling is affected in HEK293 cells lacking ESCRT-I, we assessed the phosphorylation of S6K1 kinase target, S6 protein. We observed that S6 phosphorylation is a good indicator of mTORC1 activity as treatment with INK128 compound (mTOR kinase inhibitor) caused a very strong reduction of this phosphorylation levels in control HEK293 cells (**Fig. S5A**). Importantly, ESCRT-I deficiency did not affect basal S6 phosphorylation and treating TSG101- or VPS28-depleted cells with INK128 reduced S6 phosphorylation to the same level as observed in INK128-treated control cells (**Fig. S5A**). Hence, we confirmed that in HEK293 cells, similarly to RKO cells, lack of ESCRT-I has no effect on general mTORC1 signaling.

As we have reported previously, HEK293 cells show a strong induction of TFEB/TFE3 signaling upon ESCRT-I depletion, that is a hallmark of activating the substrate-specific mTORC1 signaling due to lysosomal starvation [11]. Intriguingly, TFEB transcription factor was shown to stimulate the expression of genes involved in fatty acid oxidation [35]. Hence, to investigate any potential impact of TFEB/TFE3 signaling on regulating the transcription of amino acid or fatty acid oxidation genes upon ESCRT-I depletion, we depleted simultaneously TFEB and TFE3 using siRNA and analyzed the cells at 3 dpt. We observed that TFEB/TFE3 depletion alone led to significantly reduced mRNA levels of ALDH6A1 and BCKDHB but not of other analyzed enzymes in control cells (**Fig. S5B**). However, co-depletion of TFEB/TFE3 and ESCRT-I potentiated the reduced expression of all analyzed metabolic genes that occurs due to removal of TSG101 or VPS28 (**Fig. S5B**). Hence, we found that although TFEB/TFE3 signaling may promote the expression of genes involved in oxidative metabolism of amino acids and fatty acids, ESCRT-I depletion reduces expression of these genes despite TFEB/TFE3 signaling activation.

Collectively, we did not find evidence that the transcriptional regulation of cell metabolism observed in cells lacking ESCRT-I occurred due to the regulation of general or substrate-specific mTORC1 signaling.

### Activation of canonical NFκB signaling contributes to the reduced expression of amino acid or fatty acid oxidation genes

Having ruled out any clear contribution of mTORC1 signaling in regulation of metabolic gene expression upon ESCRT-I deficiency, we addressed a potential involvement of inflammatory/stress response pathways. Consistent with our previous study [27], ESCRT-I depletion in HEK293 cells led to the induction of canonical and non-canonical NFκB signaling, assessed by the activation of two relevant NFκB transcription factors, namely: elevated phosphorylation of RELA protein and increased levels of p52 protein (**Fig. 5A**), respectively [27, 28]. The two NFκB pathways require activation of distinct upstream kinase complexes that however share a common component, Inhibitory-κB Kinase α (IKKA) [48]. Given that the canonical, RELA-dependent, NFκB pathway may repress the expression of oxidative metabolism genes and promote glycolysis [31, 32, 49], we tested whether activation of NFκB signaling was responsible for the changes in metabolic gene expression observed upon ESCRT-I deficiency.

**Figure 5.**
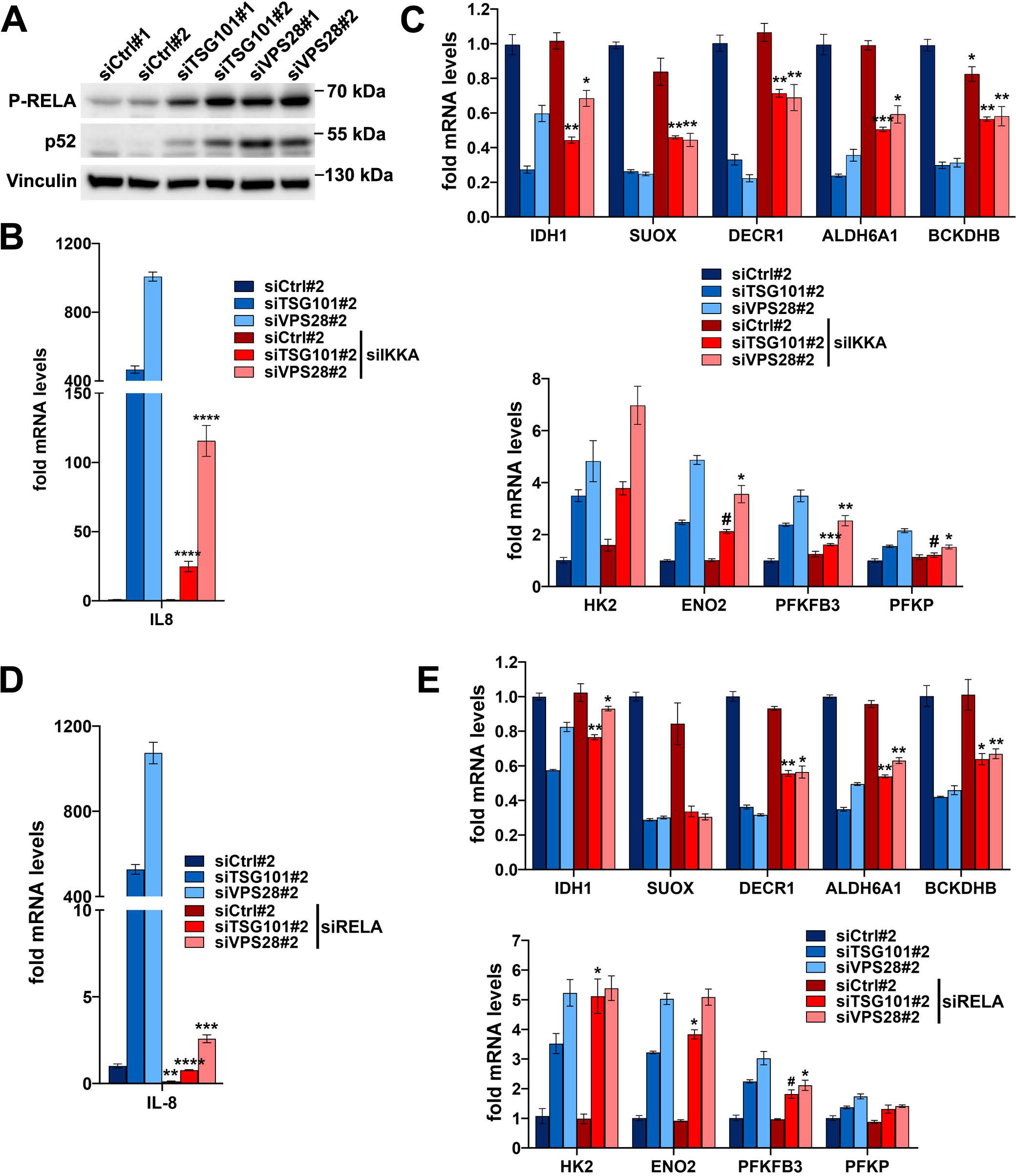
The reduced expression of genes encoding enzymes of amino acid or fatty acid oxidation in cells lacking ESCRT-I occurs in part due to activation of NFκB signaling. (A) Western blots showing the levels of phosphorylated RELA protein (P-RELA) and total p52 protein in HEK293 cells depleted of ESCRT-I using two single siRNAs for each component (siTSG101#1 or siTSG101#2, siVPS28#1 or siVPS28#2), as compared to control cells (treated with non-targeting siRNAs, Ctrl#1 or #2). The analysis was performed at three days post siRNA transfection (3 dpt) with vinculin used as a gel loading control. **(B-C)** qPCR results showing the expression of genes encoding IL-8 (in B) or indicated oxidative (top graph in C) or glycolytic enzymes (bottom graph in C) in control (siCtrl#2) or ESCRT-I- deficient (siTSG101#2 or siVPS28#2) HEK293 cells with single depletion or with co-depletion of IKKA (siIKKA) at 4 dpt. The results presented as fold changes with respect to siCtrl#2. Mean values (n = 4 ± SEM) are presented. **(D-E)** qPCR results showing the expression of genes encoding IL-8 (in D) or indicated oxidative (top graph in E) or glycolytic enzymes (bottom graph in E) in control (siCtrl#2) or ESCRT-I-deficient (siTSG101#2 or siVPS28#2) HEK293 cells with single depletion or with co-depletion of RELA (siRELA) at 4 dpt, presented as fold changes with respect to siCtrl#2. Mean values (n = 3 ± SEM) are presented. Statistical significance in (B-E) tested by comparison of cells lacking only TSG101 or VPS28 to those lacking these proteins and IKKA or RELA. ^#^P < 0.1, *P < 0.05, **P < 0.01, ***P < 0.001, ****P < 0.0001.

First, we simultaneously depleted TSG101 or VPS28 proteins with IKKA kinase to block the activation of both, the canonical and non-canonical NFκB branches upon ESCRT-I deficiency and analyzed the cells at 4 dpt. As observed by qRT-PCR analysis, we achieved efficient silencing of genes encoding these proteins, although depletion of IKKA led to upregulated mRNA levels of VPS28 and lack of ESCRT-I caused increased IKKA mRNA abundance (**Fig. S6A**). Cells lacking ESCRT-I at 4 dpt exhibited a very potently induced expression of a gene encoding the proinflammatory cytokine IL-8 (**Fig. 5B**), a transcriptional target of NFκB [50]. Its mRNA levels rose in cells lacking TSG101 by around 500-fold and in cells lacking VPS28 by roughly 1000-fold with respect to control cells (**Fig. 5B**). Removal of IKKA prevented this strong increase (**Fig. 5B**), although the elevated levels of IL-8 mRNA were still observed in cells lacking IKKA and TSG101 (by 25-fold) or VPS28 (by 120-fold with respect to control cells). Importantly, although the lack of IKKA had no effect on the expression of amino acid or fatty acid oxidation genes in control cells, it partially prevented their reduced expression due to lack of ESCRT-I (**Fig. 5C**).

To verify the involvement of only the canonical NFκB pathway in the regulation of oxidative gene expression upon ESCRT-I depletion, we inactivated this pathway in control or ESCRT-I-deficient cells by simultaneous co-depletion of RELA (**Fig. S6B**). As in the case of IKKA depletion, removal of RELA strongly prevented the elevated levels of IL-8 mRNA (**Fig. 5D**) and partially prevented the reduced mRNA levels of most oxidative enzymes observed upon TSG101 or VPS28 depletion (**Fig. 5E**).

In contrast to what we observed for amino acid or fatty acid oxidation genes, depletion of IKKA or RELA did not have a clear impact on the on the expression of genes encoding glycolytic enzymes upon ESCRT-I deficiency (**Fig. 5C and 5E**). Only the elevated expression of PFKFB3 due to TSG101 or VPS28 depletion was modestly mitigated by simultaneous IKKA (**Fig. 5C**) or RELA (**Fig. 5E**) depletion.

Cumulatively, these results showed that the activation of the canonical NFκB pathway contributes to the reduced expression of amino acid or fatty acid oxidation genes but is not responsible for the induced expression of the glycolytic metabolism genes upon ESCRT-I deficiency.

### Activation of JNK signaling contributes to induced expression of glycolytic genes in cells lacking ESCRT-I

We further reasoned that the induced expression of glycolytic genes in cells lacking ESCRT-I could occur due to activation of stress response pathways other than NFκB. Glycolysis can be stimulated by the JNK signaling [33, 51], that was found to be induced upon ESCRT inactivation [29, 30]. However, whether ESCRT dysfunction causes JNK activation in mammalian cells has not been reported. To address this, we analyzed by western blotting the effect of removing ESCRT-I on phosphorylation of JNK1/2 kinases. Although undetectable in control cells, we observed a strong JNK1/2 phosphorylation in cells lacking ESCRT-I (**Fig. 6A**).

**Figure 6.**
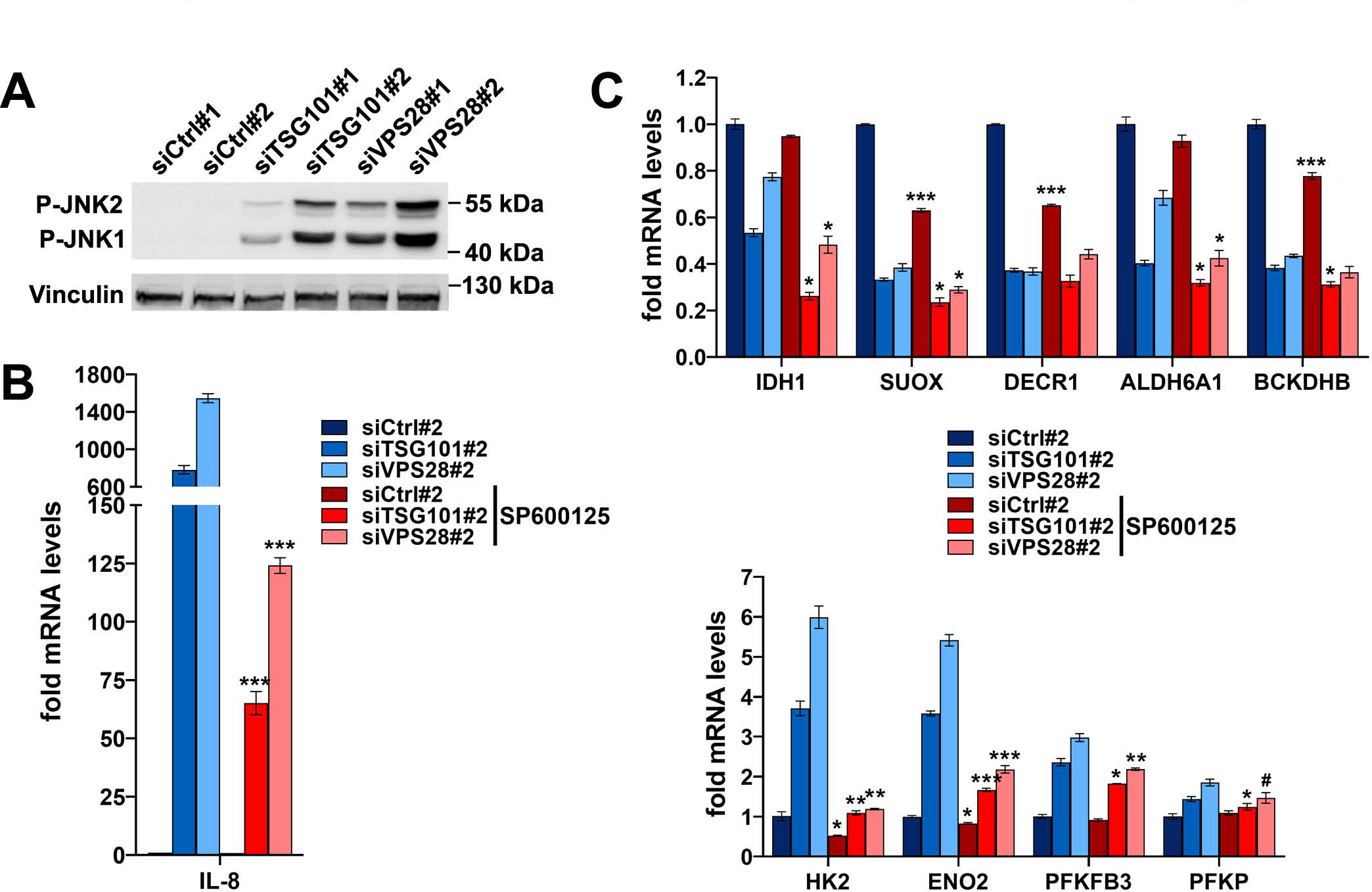
The induced expression of glycolytic metabolism genes in cells lacking ESCRT-I occurs in part due to activation of JNK signaling. (A) Western blots showing the levels of phosphorylated JNK1 and JNK2 proteins in cells depleted of ESCRT-I using two single siRNAs for each component, as compared to control cells. The analysis was performed at 3 dpt with vinculin used as a gel loading control. **(B-C)** qPCR results showing the expression of genes encoding IL-8 (in B) or the indicated oxidative (top graph in C) or glycolytic enzymes (bottom graph in C) at 4 dpt in control cells (siCtrl#2) or cells lacking ESCRT-I, upon 72 h treatment with DMSO or 50 μM SP600125 compound. Mean values (n = 3 ± SEM) are presented. Statistical significance tested by comparing the results for SP600125-treated siCtrl#2, siTSG101#2 or siVPS28#2 with results for respective DMSO-treated cells. ^#^P < 0.1, *P < 0.05, **P < 0.01, ***P < 0.001.

To investigate the involvement of JNK signaling in the transcriptional shift from oxidative to glycolytic metabolism upon ESCRT-I depletion, we treated control or ESCRT-I-deficient cells with inhibitor of JNK kinases, SP600125. This compound did not affect the depletion efficiencies of TSG101 or VPS28 (**Fig. S6C**) but strongly prevented the activated expression of the IL-8 gene (**Fig. 6B**) that in addition to NFκB is also induced by JNK signaling [52, 53]. Interestingly, SP600125 caused lower mRNA levels of SUOX, DECR1 and BCKDHB in control cells but overall did not prevent the reduced expression of the analyzed amino acid or fatty acid oxidation genes upon TSG101 or VPS28 depletion (**Fig. 6C**). However, the JNK inhibitor partially prevented the induction of glycolytic gene expression in ESCRT-I-deficient cells, having particularly strong effect on HK2 and ENO2 mRNA levels (**Fig. 6D**).

The above results showed that in cells lacking ESCRT-I the activation of JNK signaling contributes to the observed metabolic reprogramming by activating the expression of glycolytic genes.

### Inhibition of lysosomal degradation phenocopies metabolic changes observed upon ESCRT-I depletion

Induction of several signaling pathways upon ESCRT inactivation can be due to impaired endolysosomal degradation of their upstream regulators [21]. To address whether the transcriptional reprogramming towards glycolytic metabolism in cells lacking ESCRT-I could be caused by impaired lysosomal degradation, we tested the effects of inhibiting this process using bafilomycin A1 (BafA1). By qRT-PCR analysis, we observed that treatment with BafA1 led to reduced expression of all tested amino acid or fatty acid oxidation genes and increased the expression of all tested glycolytic metabolism genes (**Fig. 7A**). Next, we analyzed whether upon BafA1 treatment, these transcriptional effects are associated with induction of canonical NF-κB or JNK pathways that we found to be implicated in the metabolic shift due to ESCRT-I depletion. We observed that BafA1 strongly increased phosphorylation of RELA protein but had no effect on the levels of p52 transcription factor (**Fig. 7B**), indicating the activation of canonical but not non-canonical NF-κB signaling. Moreover, we discovered that BafA1 caused a very prominent induction of JNK1/2 phosphorylation (**Fig. 7B**).

**Figure 7.**
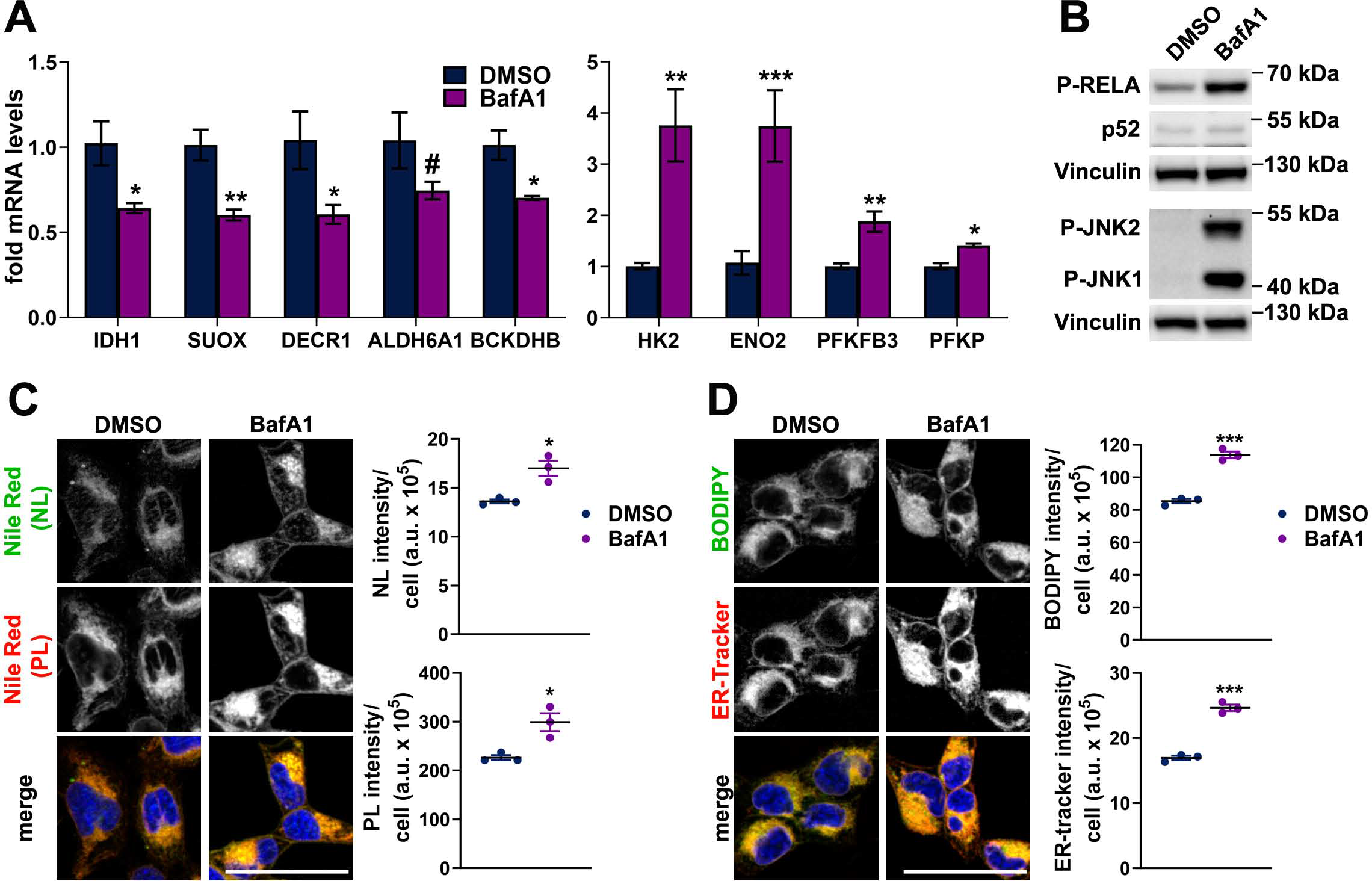
Pharmacological inhibition of lysosomal degradation recapitulates the effects of ESCRT-I depletion on metabolic gene expression and lipid accumulation. (A) qPCR results showing the expression of genes encoding the indicated oxidative (left) or glycolytic (right) enzymes in HEK293 cells treated for 24 h with DMSO or 20 nM Bafilomycin A1 (BafA1). Mean values (n = 4 ± SEM) are presented. Statistical significance tested by comparison to cells treated with DMSO. **(B)** Western blots showing the levels of phosphorylated RELA, JNK1 and JNK2 proteins and levels of total p52 protein in HEK293 cells treated for 24 h with DMSO or 20 nM BafA1. Vinculin used as a gel loading control. **(C-D)** Maximum intensity projection confocal images, showing the intracellular distribution of neutral lipids – NL (green), or phospholipids – PL (red) stained with Nile Red dye (shown in C) as well as the intracellular distribution of NLs stained with BODIPY 493/503 and the ER stained with ER-tracker Red (green or red, respectively in D) in live cells treated for 24 h with DMSO or 20 nM BafA1. Cell nuclei marked with Hoechst stain (blue). Scale bar, 50 μm. The dot plots on the right show total fluorescence intensities per cell (expressed in arbitrary units, a.u.). Values derived from independent experiments (dots) and their means (n = 3 ± SEM) are presented. ^#^P < 0.1, *P < 0.05, **P < 0.01, ***P < 0.001.

As we observed that BafA1 treatment recapitulated the effects of ESCRT-I depletion on metabolic gene expression and activation of canonical NF-κB and JNK signaling, we investigated whether it also affected lipid homeostasis. By confocal microscopy of HEK293 cells stained with NR or BODIPY, we observed that BafA1 led to intracellular accumulation of NLs and PLs (**Fig. 7C-D**). Moreover, it also led to elevated ER content (**Fig. 7D**). Hence, pharmacological inhibition of lysosomal degradation phenocopied the effects of ESCRT-I depletion on the expression of genes involved in nutrient metabolism as well as intracellular lipid content. This suggests that impaired lysosomal degradation could underly the transcriptional reprogramming of cell metabolism upon ESCRT-I depletion.

## DISCUSSION

Collectively, our results show that functional ESCRT-I promotes oxidative metabolism of amino acids and fatty acids over glycolysis by restricting stress and inflammatory pathways, potentially due to involvement of this complex in delivering cargo for lysosomal degradation. A metabolic switch towards aerobic glycolysis occurs during many important cellular processes, such as somatic cell reprogramming, macrophage polarization, tissue remodeling during inflammation and carcinogenesis [8, 54–56] but still many of its underlying aspects are not well understood. A growing number of studies has shown that endocytic trafficking affects cell metabolism [57, 58] and that particular endocytic machineries may regulate oxidative or glycolytic metabolism [59–61]. Hence, it is likely that endocytic proteins are involved in metabolic reprogramming events.

By transcriptomic analysis and its validation, we discovered that removal of ESCRT-I in HEK293 or HepG2 cells leads to opposite regulation of the expression of two particular groups of metabolism-related genes: those involved in oxidative metabolism of carboxylic acid-containing nutrients (with downregulated expression) and those related to glycolysis (with upregulated expression). The group of downregulated genes includes those encoding enzymes that mediate initial reactions of oxidative breakdown of FFAs and BCAAs. These reactions provide esters of CoA and electron carriers, such as NADH, thus molecules that are incorporated into further oxidation in mitochondria within the citric acid cycle and oxidative phosphorylation [2, 62, 63]. Hence, although the metabolism-related genes, whose expression is reduced upon ESCRT-I depletion, do not encode key mitochondrial components, they function in obtaining energy from oxidation of specific nutrients, such as FFAs and BCAAs.

The group of upregulated genes implicated in glycolysis, although small, encompasses those encoding rate-limiting glycolytic enzymes, HK2, ENO2 and PFK, as well as an enzyme that potently stimulates glycolysis, PFKFB3 [64, 65]. Consistently with upregulated expression of these genes, we discovered that cells lacking ESCRT-I have elevated lactate production and increased glycolysis-related ECAR, hence they perform aerobic glycolysis [1]. We propose that changes in metabolic gene expression observed upon ESCRT-I deficiency may lead to preferential usage of glucose over FFAs and BCAAs for energy production. However, as the ATP-linked respiration of cells lacking ESCRT-I is not reduced, it is possible that the glycolysis-derived pyruvate in these cells is used not only for lactate production but also enters the mitochondria to generate acetyl-CoA fueling mitochondrial respiration.

The here reported reduced expression of amino acid and fatty acid oxidation genes upon ESCRT-I dysfunction coincides with a broad accumulation of intracellular membranes. As we described previously, cells lacking ESCRT-I have enlarged endosomes and lysosomes, that are hallmarks of inhibited endolysosomal degradation [11, 27]. In the current study we also show that these cells accumulate mitochondria and lipid droplets, and their ER is expanded. The increased abundance of mitochondria and the enlarged ER in cells lacking TSG101 or VPS28 are likely due to impaired autophagic degradation of these organelles [23, 66]. Accumulation of lipid droplets could be a result of diminished lipid transport from the droplets into mitochondria proposed to be mediated by ESCRT-I proteins [67]. The profoundly impaired turnover of organelles and intracellular membranes in the absence of ESCRT-I likely results in a reduced supply of molecules that are recycled from lysosomal degradation, including FFAs. Our finding that cells lacking ESCRT-I exhibit not reduced but even slightly elevated FFA levels allows hypothesizing that transcriptional regulation of genes involved in FFA metabolism and synthesis in these cells compensates for reduced fatty acid delivery from lysosomes. However, further experiments should address the rate of fatty acid oxidation upon ESCRT-I deficiency.

In this study, we corroborated our previous observation that lack of ESCRT-I does not affect the general mTORC1 signaling [11]. Therefore, we reasoned that the observed changes in the expression of metabolic genes are not due to general regulation of mTOR kinase activity. However, we were intrigued by finding that the reduced expression of genes related to oxidative metabolism occurs in cells in which we previously observed a prominent activation of TFEB/TFE3 transcription factors [11] that may promote expression of genes related to fatty acid oxidation [35]. By addressing this subject, we clarified that the activation of TFEB/TFE3 transcription factors partially prevents the reduced expression of genes related to fatty acid and amino acid oxidation. The activation of TFEB/TFE3 factors serves as a homeostatic response in an attempt to rescue the impaired lysosomal degradation [68]. Hence, stronger effect on the expression of amino acid and fatty acid oxidation genes upon ESCRT deficiency in cells lacking TFEB/TFE3 is consistent with our interpretation that the changes in metabolic gene expression in cells lacking ESCRT-I are due to inhibited lysosomal degradation.

Importantly, we demonstrate that activation of the NFκB pathway in mammalian cells lacking ESCRT-I, in addition to promoting inflammatory signaling, inhibits the expression of genes encoding enzymes of amino acid or fatty acid oxidation. This effect may be indirect, e.g. due to NFκB-dependent regulation of other transcription factors as NFκB signaling may affect transcriptional regulators of metabolism such as PPAR nuclear receptors or SIRT1 deacetylase [31, 49]. The role of the NFκB signaling in regulation of glycolytic metabolism is not well established. Some reports suggest that it promotes [32, 69] and some others that it inhibits glycolysis [70, 71]. We observe that preventing the activation of NFκB signaling does not interfere with increased expression of glycolytic genes upon ESCRT-I deficiency.

Our findings that ESCRT-I deficiency in mammalian cells leads to activation of the JNK pathway extend similar observations made previously for mutants of genes encoding ESCRT-0 or ESCRT-II components [29, 30]. Whereas those studies showed that the activation of JNK signaling may affect cell proliferation or survival upon ESCRT dysfunction [29, 30], we demonstrate that activation of this signaling participates in inducing the expression of glycolytic genes in cells lacking TSG101 or VPS28. Intriguingly, the exact three glycolytic genes with the highest upregulation upon ESCRT-I deficiency (encoding HK2, ENO2 and PFKFB3) were reported to have increased expression in cancer-associated fibroblasts due to activation of JNK signaling upon compression-induced stress [51]. Hence, these genes appear to be a part of a particular JNK-dependent transcriptional response commonly activated upon various stress conditions. Further investigation is required to unravel which transcription factors mediate this response downstream of JNK kinases in cells lacking ESCRT-I.

Although it has been previously proposed that pharmacological inhibition of lysosomal degradation may promote glycolytic metabolism [72], to our knowledge the effects of such treatment on the expression of genes related to glycolytic or oxidative metabolism has not been demonstrated. Here we show that BafA1 causes a transcriptional reprogramming from amino acid and fatty acid oxidation to glycolytic metabolism, similar to what we observe in cells lacking ESCRT-I. It is likely that impaired lysosomal degradation of membrane receptors activates stress and inflammatory signaling pathways to mediate the transcriptional shift occurring upon deficiency of ESCRT-I or treatment with BafA1, to be formally verified in the future.

Overall, the results of our study suggest that ESCRT-I activity could potentially promote oxidative metabolism, while its inactivation could favor glycolysis. Further studies should address whether the abundance and/or activity of ESCRT-I is regulated upon stem cell differentiation that involves reprogramming from glycolysis to oxidative metabolism [73–75], or upon inducing cell pluripotency, inflammation or oncogenesis that involve a reprogramming from oxidative to glycolytic metabolism [8, 54–56].

## MATERIALS AND METHODS

### Antibodies

The following antibodies were used: anti-TSG101 (Cat# ab83), anti-VPS28 (Cat# ab167172) and anti- SUOX (Cat# ab129094) from Abcam; anti-P-RELA (Cat# 3033), anti-p52 (Cat# 4882), anti-P-S6 (Cat# 2211) and anti-P-JNK1/2 (Cat# 9255) from Cell Signaling Technologies; anit-IDH1 (Cat# GT1521) from GeneTex; anti-ALDH6A1 (Cat# sc-271582) and anti-HK2 (Cat# sc-374091) from Santa Cruz; anti-PFKFB3 (Cat# 13763-1-AP) from Proteintech; anti-vinculin (Cat# V9131-2ML) from Sigma-Aldrich; secondary horseradish peroxidase (HRP)-conjugated goat anti-mouse and goat anti-rabbit from Thermo Fisher Scientific.

### Plasmids

pLenti-CMV-MCS-GFP-SV-puro (Addgene plasmid #73582) was a gift from Katarzyna Mleczko-Sanecka. psPAX2 (Addgene plasmid #12260) and pMD2.G (Addgene plasmid #12259) lentiviral packaging plasmids were a gift from Didier Trono.

### Cell culture and treatment

HEK293 and HEK293T embryonic kidney cells were maintained in Dulbecco’s modified Eagle’s medium (DMEM, Sigma-Aldrich, M2279). HepG2 hepatoblastoma cells were cultured in Eagle’s minimum essential medium (EMEM, ATCC, 30-2003). Both medium types were supplemented with 10% (v/v) fetal bovine serum (FBS, Sigma-Aldrich, F7524) and 2 mM L-Glutamine (Sigma-Aldrich, G7513). All cell lines were regularly tested as mycoplasma-negative and their identities were confirmed by short tandem repeat (STR) profiling performed by the ATCC Cell Authentication Service.

Bafilomycin A1 (Sigma-Aldrich, B1793) at 20 nM concentration was applied for 24 h to inhibit lysosomal degradation. To inhibit JNK activity, SP600125 was used at 50 μM concentration for 72 h. To inhibit mTOR kinase activity INK128 was used at 100 nM concentration for 48 h.

### Cell transfection and lentiviral transduction

HEK293 cells were seeded on 6-well (1x10^5^ cells/well) or 12-well (1x10^5^ cells/well) plates for western blotting and quantitative real-time PCR (qRT-PCR) experiments or on 0.2% gelatin (Sigma Aldrich, G1890)-covered 96-well plate (Grainer Bio-One, 655-090) (2.5x10^3^ cells/well) for confocal microscopy. HepG2 cells were seeded on 6-well plates (2x10^5^ cells/well) for western blotting or on gelatin-covered 96-well plates (4x10^3^ cells/well) for microscopy. 24 h after seeding, cells were transfected with 20 nM siRNAs using Lipofectamine™ RNAiMAX Transfection Reagent (Thermo Fisher Scientific, 13778150) according the manufacturer’s instructions and imaged or harvested after 3 or 4 days post transfection (dpt). The following Ambion Silencer Select siRNAs (Thermo Fisher Scientific) were used: Negative Control No. 1 (siCtrl#1, 4390843) Negative Control No. 2 (4390846) and Negative Control No. 3 (siCtrl#3, custom-made, UACGACCGGUCUAUCGUAG); siTsg101#1 (s14439), siTsg101#2 (s14440), siVps28#1 (s27577), siVps28#2 (s27579), siTFEB (s15496), siTFE3 (s14031), siIKKA (s3076), siRELA (s11916). In experiments with simultaneous knockdown of two genes, the total concentration of siRNA was adjusted to 40 nM using siCtrl#2.

For CRISPR/Cas9-mediated knock-out of the gene encoding TSG101, 2 control non-targeting gRNA sequences and 1 targeting gRNA sequence [76] were cloned into the lentiCRISPRv2 vector. Lentiviral particles were produced in HEK293T cells using packaging plasmids: psPAX2 and pMD2.G, as described elsewhere [77]. Subsequently, 1x10^6^ HepG2 cells were grown in 5 ml of virus-containing EMEM medium on P60 dish for 1 h. After the infection, the cells were kept in fresh EMEM for 24h. Then, cells were split and grown in selection EMEM medium containing 1 µg/ml puromycin for 7 days. The sequences of gRNAs used in this study are: gCtrl#1- CGCTTCCGCGGCCCGTTCAA, gCtrl#2- CTGAAAAAGGAAGGAGTTGA, gTsg101#1- AGGGAACTAATGAACCTCAC.

### Microarray analysis

HEK293 cells were transfected with control (siCtrl#1 or siCtrl#2) or ESCRT-I-targeting (siTSG101#1, siTSG101#2, siVPS28#1 or siVps28#2) siRNAs in three biological repetitions. Three days post transfection (3 dpt) cells were collected and cRNA was prepared as described elsewhere [78]. cRNA samples hybridized to the Human HT-12 BeadChip array (Illumina) were scanned on the BeadArray Reader (Illumina) and analyzed using BeadStudio v3.0 (Illumina) and BeadArray R package v1.10.0 software according to a procedure described elsewhere [26]. Upon quantile normalization, data were log2 transformed and statistical analysis of the results was performed using a t-test followed by false discovery rate (FDR) correction for multiple testing according to the Benjamini and Hochberg method.

Ingenuity Pathway Analysis software was used to identify canonical pathways potentially activated or inhibited based on the common list of genes whose expression was significantly increased or decreased (FDR<0.05) in cells lacking TSG101 or VPS28. GO analysis of biological processes or cellular components was performed using enrichGO function from clusterProfiler R-package (version 4.10.0; [79]), on the list of the genes with commonly downregulated expression (FDR<0.05, log2 fold change to controls ≤ - 0.6). Heatmaps were plotted using ComplexHeatmap (version 2.16.0, [80]). The visualizations were performed in R version 4.3.1 (https://www.R-project.org). Heatmaps for genes encoding enzymes of glucose catabolism and proteins involved in biosynthesis of FFAs, TGs or PLs visualize the expression of genes from Gene Ontology lists (glycolytic process_GO:0006096, acetyl−CoA biosynthetic process from pyruvate (GO:0006086), fatty acid biosynthetic process_GO:0006633, triglyceride biosynthetic process_GO:0019432, glycerophospholipid biosynthetic process_GO:0046474) that were manually curated to remove unrelated genes.

### Quantitative real-time PCR (qRT-PCR)

Total RNA was isolated from cells using High Pure Isolation Kit (Roche, 11828665001) according to the manufacturer’s instruction. For cDNA preparation, 1000 ng of total RNA, random nonamers (Sigma-Aldrich, R7647), oligo(dT)23 (Sigma-Aldrich, O4387) and M-MLV reverse transcriptase (Sigma-Aldrich, M1302) were used. In most qRT-PCR analyzes, the NCBI Primer designing tool was used to design primers that were custom-synthesized by Sigma-Aldrich (primer sequences listed in **Table 1**) and cDNA sample amplification was performed with the KAPA SYBR FAST qPCR Kit (KapaBiosystems, KK4618). Only to analyze the expression of amino acid or fatty acid oxidation genes at 3 days post transfection (shown in Fig. 1E), TaqMan™ assays (ThermoFisher Scientific, 4331182) with TaqMan™ Gene Expression Master Mix (ThermoFisher Scientific, 4369016) were used (assays listed in **Table. 2**). In both, using custom-synthesized primers and TaqMan™ assays, the 7900HT Fast Real-Time PCR thermocycler (Applied Biosystems) was used for DNA amplification. Obtained data were normalized according to the expression level of the housekeeping genes encoding ACTB (β-actin) or HPRT1 (Hypoxanthine Phosphoribosyltransferase 1) proteins. Results are presented as fold changes compared to the average values obtained for control cells or to values obtained using siCtrl#2 (in co-depletion experiments).

**Table 1.**
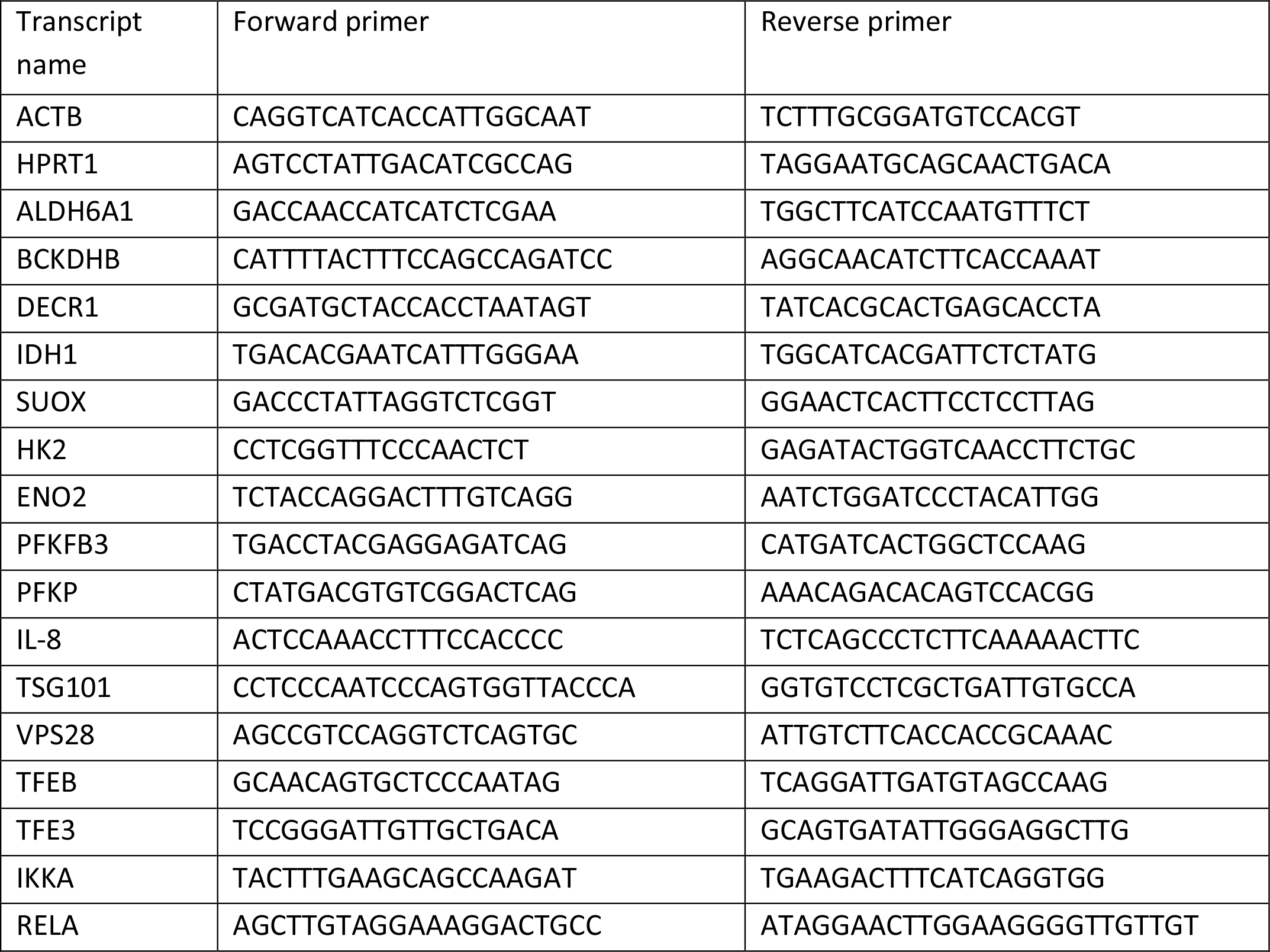
Sequences of primers used for qRT-PCR

**Table 2.**
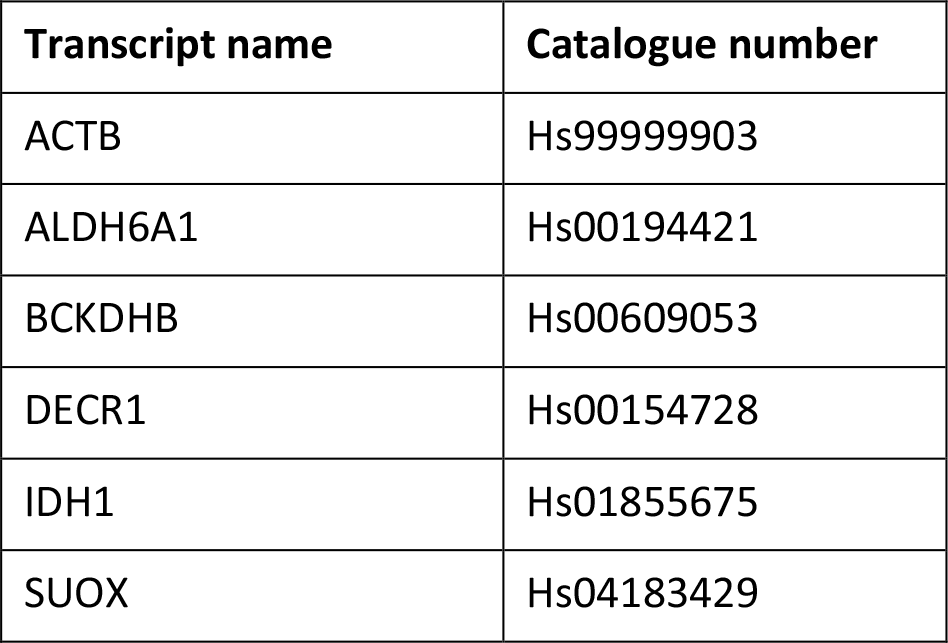
TaqMan assays used for qRT-PCR

### Western blotting

Cells were lysed in RIPA buffer (1% Triton X-100, 0.5% sodium deoxycholate, 0.1% SDS, 50 mM Tris pH 7.4, 150 mM NaCl, 0.5 mM EDTA) supplemented with phosphatase inhibitor cocktails 2 and 3 (Sigma-Aldrich, P5726 and P0044) and protease inhibitor cocktail (6 μg/ml chymostatin, 0.5 μg/ml leupeptin, 10 μg/ml antipain, 2 μg/ml aprotinin, 0.7 μg/ml pepstatin A and 10 μg/ml 4- amidinophenylmethanesulfonyl fluoride hydrochloride; Sigma-Aldrich). BCA Protein Assay Kit (Thermo Fisher Scientific, 23225) was used to measure protein concentration. Subsequently, 15-25 µg of total protein per sample were resolved on 8%, 10% or 12% SDS-PAGE and transferred onto nitrocellulose membrane (Amersham Hybond, GE Healthcare Life Science, 10600002). Membranes were blocked in 5% milk PBS with 0.1% Tween followed by incubation with specific primary and secondary antibodies. For signal detection, Clarity Western ECL Substrate (BioRad, 170-5061) and ChemiDoc imaging system (Bio-Rad) were applied. Image Lab 6.0.1 software (Bio-Rad) was used for densitometric analysis of western blotting bands. The raw data were normalized to vinculin or tubulin band intensities and presented as fold levels to the average values obtained for control cells.

### Lipid extraction, separation and gas chromatography-mass spectrometry (GC-MS) analysis

HEK293 cells were cultured in 100 mm dishes and 3 days post siRNA transfection were harvested washed with PBS and cell pellets were processed for lipid extraction. Lipids were extracted according to the Bligh and Dyer method [81]. Each sample was homogenized in a glass tube in chloroform:methanol (2:1) containing 0.01% (w/v) butylated hydroxytoluene (BHT). Then distilled water was added to each tube and the tubes were vortexed and centrifugated at 3,000 rpm for 10 minutes at 4°C. After centrifugation, the resulting two-phase system was separated and the lower phase containing lipids was collected.

Lipid extracts were separated by thin-layer chromatography on silica gel 60 plates (Merck, Darmstadt, Germany) in heptane/isopropyl ether/glacial acetic acid (60/40/4, vol/vol/vol) with authentic standards. For fatty acid analysis, bands corresponding to free fatty acids (FFAs) triglycerides (TGs), diglycerides (DAGs), and phospholipids (PLs) were visualized with 0.2% 2,3-dichlorofluorescein, scraped from the plate into screw-capped glass tubes and transmethylated in the presence of 14% boron trifluoride in methanol. The resulting FA methyl esters were extracted with hexane and subjected to Agilent 7890A-5975C GC-MS instrument with an Agilent 19091N-205 capillary column (Agilent Technologies, Santa Clara, CA, USA). Peak alignment, purity and quality analyses were performed using Agilent MSD Productivity ChemStation software. Methyl nonadecanoate was used as an internal standard for further compound quantification by selected ion monitoring. For normalization, the measured concentrations of various lipid classes (nmol/μL) were divided by protein concentration (μg/μL) that was assessed using BCA Protein Assay Kit (Thermo Fisher Scientific, 23225).

### Live cell staining and microscopy

For live cell imaging, cells grown on 0.2% gelatin-coated 96-well plates (Greiner Bio-One, 655-090) were incubated for 15 min with 1 μg/mL Nile Red (N1142), 0.4 μg/mL BODIPY 493/503 (D3922), 1 μM ER-tracker Red (E34250), 100 nM MitoTracker™ Deep Red FM (M22426), 100 nM TMRE (T669) or 10 μg/mL Hoechst 33342 (Thermo Fisher Scientific, H3570) and imaged immediately. Plates were scanned using Opera Phenix high content screening microscope (PerkinElmer) with 40 × 1.1 NA water immersion objective. Harmony 4.9 software (PerkinElmer) was applied for image acquisition and their quantitative analysis. For quantification of integral fluorescence intensity per cell, more than 10 microscopic fields were analyzed per each experimental condition. Maximum intensity projection images were obtained from 3 z-stack planes with 1 μm interval. Pictures were assembled in ImageJ and Photoshop (Adobe) with only linear adjustments of contrast and brightness.

### Lactate content analysis

Colorimetric analysis of intracellular abundance of lactate was performed using Lactate Assay Kit (Sigma-Aldrich, MAK064) according the manufacturer’s instructions. For this analysis, HEK293 cells were cultured in 12-well plates and 4 days post transfection with siRNAs (4dpt) they were harvested into assay buffer and homogenized using a tissue homogenizer. For normalization, the measured amount of lactate in the assay buffer (nmol/μL) was divided by DNA concentration (μg/μL) that was assessed using NanoDrop 1000 spectrophotometer (Thermo Fisher Scientific).

### Energy metabolism analysis

Oxygen consumption rate (OCR) as well as extracellular acidification rate (ECAR) were measured with the Seahorse XFe24 Analyzer (Agilent Technologies, Santa Clara, CA, USA). To achieve this, 4x10^4^ or 8x10^4^ cells per well were plated in Seahorse 24-well cell plates for 3dpt (OCR) and 4dpt (ECAR) measurements, respectively. On the next day, cells were transfected with siRNAs at 20 nM concentration. OCR and ECAR were measured according to manufacturer’s instructions by sequential injections of compounds that modulate mitochondrial function: 4 μM oligomycin (Sigma-Aldrich) that inhibits ATP-linked respiration, 2 μM carbonyl cyanide-p-trifluoromethoxyphenylhydrazone (FCCP, from Sigma-Aldrich) that allows for maximal electron flow through the electron transport chain, thus maximal oxygen consumption, and 2 μM antimycin A (Sigma-Aldrich) together with 2 μM rotenone (Sigma-Aldrich) that shut down mitochondrial respiration. To assess glycolysis-related ECAR an additional injection of 100 mM 2-deoxy-d-glucose (2-DG, from Sigma-Aldrich), an inhibitor of glycolysis, was performed. The results (pmol of consumed oxygen per minute and mpH altered per minute) were normalized to the fluorescence intensity of 10 μM Hoechst 33342 (Thermo Fisher), added to cells during the assay, that stains DNA and reflects cell number.

### Statistical analysis

Data are shown as mean ± SEM from at least three independent biological experiments. Statistical analyses were performed with the Prism 10.0.3 (GraphPad Software). In most of the analyses, the unpaired two-tailed t test was used, with the exception of confocal microscopy and GC-MS experiments in which the ratio paired t test was used. The significance of mean comparison is indicated with ^#^P<0.1, *P < 0.05, **P < 0.01, ***P < 0.001, ****P < 0.0001. Non-significant results (P>0.1) were not indicated.

## STATEMENTS & DECLARATIONS

## Acknowledgements and funding

We are grateful to Jacek Jaworski and Katarzyna Mleczko-Sanecka for providing reagents. The work was supported by the HOMING grant (POIR.04.04.00-00-1C54/16-00) to JC and the TEAM grant (POIR.04.04.00-00-20CE/16–00) to MMiaczynska, both from the Foundation for Polish Science co-financed by the European Union under the European Regional Development Fund.

## Author contributions

The research was conceived by JC and MMiaczynska. Funding was acquired by JC and MMiaczynska. Experiments were designed by JC, and most of them were performed by JC, MW, MMazur and BJ.

Transcriptomic analysis was performed by MK and its visualization was performed by MW and MR. Analysis of Extracellular Flux using Seahorse XF24 was performed by SW and supervised by AZ. GC-MS analysis was performed by ADziewulska and supervised by ADobrzyn. The manuscript was written by JC, and edited by MMiaczynska, MR and AZ. JC and MMiaczynska supervised the work. All authors approved the manuscript.

## Data Availability

The data from microarray analysis have been deposited to ZENODO: DOI 10.5281/zenodo.10784726 (https://zenodo.org/records/10784727).

## Ethics approval

Not applicable

## Consent for publication

Not applicable

---

**Table S1.** List of genes downregulated upon ESCRT-I depletion annotated to small molecule metabolic process. Indicated are average fold changes, false discovery rates (FDR) and information whether each gene is involved in amino acid or fatty acid oxidation or in other process category.

| **GO:0044281: small molecule metabolic process** |  |  |  |
| --- | --- | --- | --- |
| **gene** | **average log2 fold change** | **FDR** | **category** |
| NAALAD2 | -1,1465 | 0,0007 | amino acid oxidation |
| SUOX | -1,1192 | 0,0318 | amino acid oxidation |
| NME3 | -1,1189 | 0,0079 | other |
| COQ5 | -1,0477 | 0,0208 | other |
| OARD1 | -1,0106 | 0,0013 | other |
| CACNA1H | -0,9718 | 0,0053 | other |
| HOGA1 | -0,9714 | 0,0028 | amino acid oxidation |
| TM7SF2 | -0,9179 | 0,0107 | other |
| IDH1 | -0,9109 | 0,0076 | fatty acid oxidation |
| EPHX2 | -0,8995 | 0,0065 | fatty acid oxidation |
| ALDH3A2 | -0,8976 | 0,0050 | fatty acid oxidation |
| IDNK | -0,8878 | 0,0121 | other |
| CLYBL | -0,8567 | 0,0121 | other |
| NUDT7 | -0,8554 | 0,0037 | fatty acid oxidation |
| ACCS | -0,8552 | 0,0330 | other |
| CYP4V2 | -0,8495 | 0,0165 | other |
| GSTM2 | -0,8487 | 0,0494 | other |
| BPHL | -0,8487 | 0,0147 | other |
| SIRT5 | -0,7967 | 0,0002 | other |
| MIDN | -0,7699 | 0,0085 | other |
| PLCB1 | -0,7625 | 0,0123 | fatty acid oxidation |
| NNT | -0,7586 | 0,0141 | other |
| PEX2 | -0,7578 | 0,0054 | fatty acid oxidation |
| ACACB | -0,7557 | 0,0336 | fatty acid oxidation |
| UQCR10 | -0,7429 | 0,0173 | other |
| SUCLG2 | -0,7360 | 0,0097 | other |
| ATP2B4 | -0,7351 | 0,0439 | amino acid oxidation |
| NR1H3 | -0,7113 | 0,0114 | other |
| BCKDHB | -0,7098 | 0,0013 | amino acid oxidation |
| ALDH7A1 | -0,7096 | 0,0015 | amino acid oxidation |
| ATP6V1B1 | -0,7087 | 0,0139 | other |
| HMGCL | -0,7057 | 0,0040 | amino acid oxidation |
| CPT2 | -0,6978 | 0,0127 | fatty acid oxidation |
| EXTL2 | -0,6928 | 0,0057 | other |
| DECR1 | -0,6877 | 0,0182 | fatty acid oxidation |
| PEX7 | -0,6849 | 0,0027 | fatty acid oxidation |
| CSAD | -0,6772 | 0,0028 | amino acid oxidation |
| GALM | -0,6737 | 0,0010 | other |
| LDHC | -0,6736 | 0,0035 | other |
| TRERF1 | -0,6684 | 0,0094 | fatty acid oxidation |
| THEM4 | -0,6629 | 0,0173 | fatty acid oxidation |
| TST | -0,6583 | 0,0049 | amino acid oxidation |
| MST1 | -0,6501 | 0,0215 | other |
| RDH10 | -0,6488 | 0,0070 | other |
| ECI2 | -0,6340 | 0,0005 | fatty acid oxidation |
| DCAKD | -0,6336 | 0,0053 | other |
| CAV1 | -0,6276 | 0,0300 | other |
| CRYZL1 | -0,6245 | 0,0334 | other |
| ALDH6A1 | -0,6178 | 0,0154 | amino acid oxidation |
| CAT | -0,6172 | 0,0294 | other |
| PPA2 | -0,6171 | 0,0068 | other |
| ACSF2 | -0,6143 | 0,0083 | fatty acid oxidation |
| UROS | -0,6115 | 0,0166 | other |
| DDIT4 | -0,6052 | 0,0315 | other |
| NUDT1 | -0,6044 | 0,0101 | other |
| PLA2G4B | -0,6028 | 0,0049 | fatty acid oxidation |
| EHHADH | -0,6022 | 0,0042 | fatty acid oxidation |

**Table S2.** List of genes downregulated upon ESCRT-I depletion annotated to mitochondrion. Indicated are average fold changes, false discovery rates (FDR) and information whether each gene is involved in amino acid or fatty acid oxidation or in other process category.

| GO:0005739: mitochondrion |  |  |  |
| --- | --- | --- | --- |
| gene | average log2 fold change | FDR | category |
| SUOX | -1,1192 | 0,0318 | amino acid oxidation |
| MSRB2 | -1,0815 | 0,0033 | other |
| COQ5 | -1,0477 | 0,0208 | other |
| NRP1 | -1,0194 | 0,0028 | other |
| BMF | -0,9904 | 0,0103 | other |
| HOGA1 | -0,9714 | 0,0028 | amino acid oxidation |
| ASB9 | -0,9615 | 0,0180 | other |
| IDH1 | -0,9109 | 0,0076 | fatty acid oxidation |
| CLYBL | -0,8567 | 0,0121 | other |
| BPHL | -0,8487 | 0,0147 | other |
| CA5B | -0,8263 | 0,0249 | other |
| CMC4 | -0,8254 | 0,0107 | other |
| IFI27L1 | -0,8239 | 0,0130 | other |
| PYROXD2 | -0,8212 | 0,0084 | other |
| SIRT5 | -0,7967 | 0,0002 | other |
| NNT | -0,7586 | 0,0141 | other |
| ACACB | -0,7557 | 0,0336 | fatty acid oxidation |
| UQCR10 | -0,7429 | 0,0173 | other |
| SUCLG2 | -0,7360 | 0,0097 | other |
| EXD2 | -0,7195 | 0,0149 | other |
| LIPT2 | -0,7169 | 0,0062 | other |
| BCKDHB | -0,7098 | 0,0013 | amino acid oxidation |
| ALDH7A1 | -0,7096 | 0,0015 | amino acid oxidation |
| HMGCL | -0,7057 | 0,0040 | amino acid oxidation |
| TMEM143 | -0,6984 | 0,0365 | other |
| CPT2 | -0,6978 | 0,0127 | fatty acid oxidation |
| DECR1 | -0,6877 | 0,0182 | fatty acid oxidation |
| SLC25A42 | -0,6787 | 0,0052 | other |
| LDHC | -0,6736 | 0,0035 | other |
| THEM4 | -0,6629 | 0,0173 | other |
| TST | -0,6583 | 0,0049 | amino acid oxidation |
| NFU1 | -0,6542 | 0,0068 | other |
| COA5 | -0,6529 | 0,0479 | other |
| TMEM135 | -0,6374 | 0,0381 | other |
| ECI2 | -0,6340 | 0,0005 | fatty acid oxidation |
| PTCD2 | -0,6303 | 0,0090 | other |
| TXNRD2 | -0,6178 | 0,0028 | other |
| ALDH6A1 | -0,6178 | 0,0154 | amino acid oxidation |
| CAT | -0,6172 | 0,0294 | other |
| PPA2 | -0,6171 | 0,0068 | other |
| ACSF2 | -0,6143 | 0,0083 | fatty acid oxidation |
| LETMD1 | -0,6123 | 0,0048 | other |
| UROS | -0,6115 | 0,0166 | other |
| BRD8 | -0,6099 | 0,0080 | other |
| PRIMPOL | -0,6067 | 0,0080 | other |
| RMDN1 | -0,6054 | 0,0163 | other |
| DDIT4 | -0,6052 | 0,0315 | other |
| NUDT1 | -0,6044 | 0,0101 | other |
| PLA2G4B | -0,6028 | 0,0049 | fatty acid oxidation |

**Figure S1.**
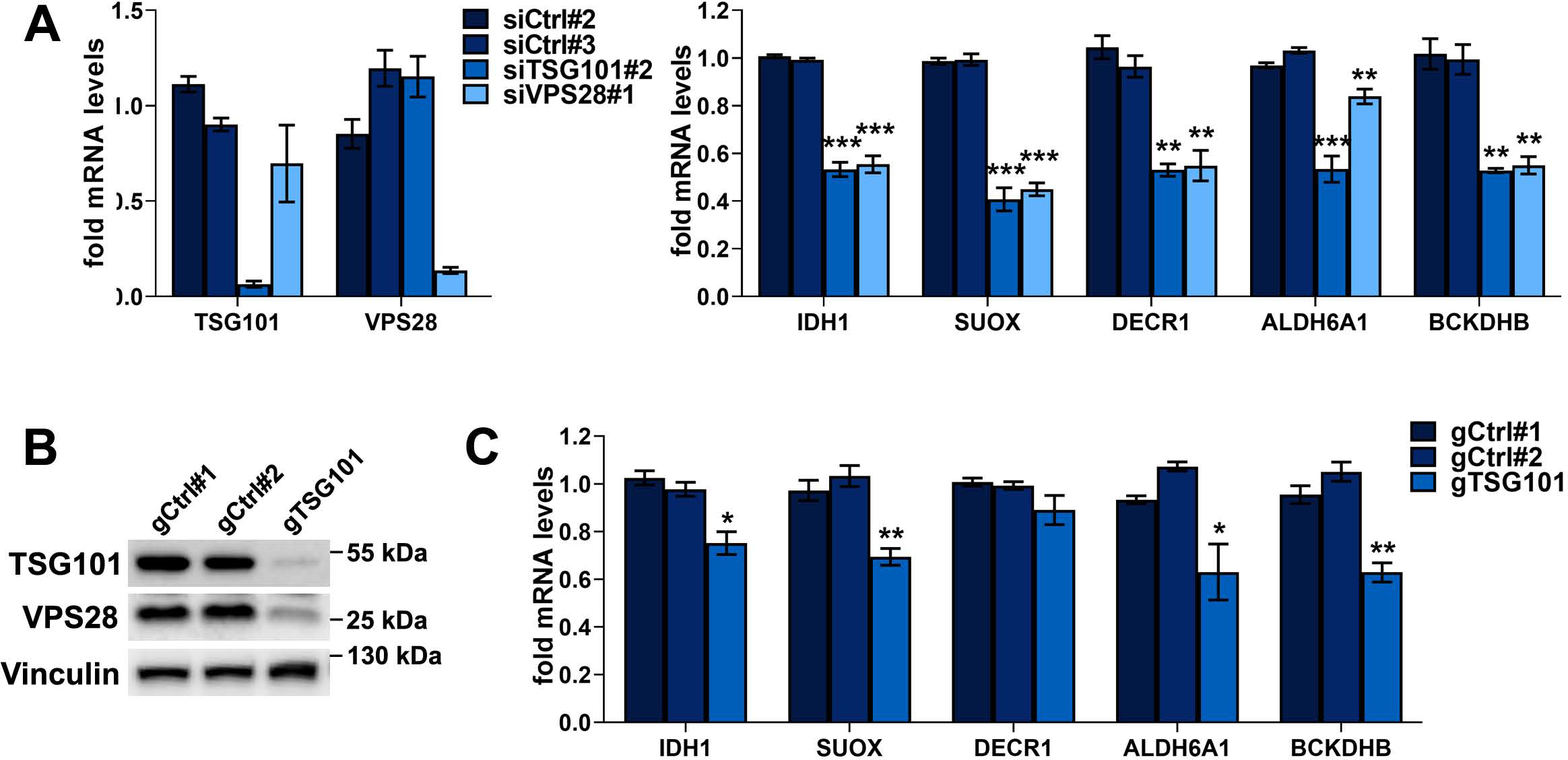
ESCRT-I dysfunction leads to reduced expression of genes involved in oxidative metabolism of amino acids and fatty acids in HepG2 cells. (A) qPCR results showing the expression of genes encoding TSG101, VPS28 and the indicated oxidative metabolism enzymes in HepG2 cells with removal of ESCRT-I using single siRNA for each component (siTSG101#2, siVPS28#1), as compared to control cells (treated with non-targeting siRNAs, Ctrl#2 or #3) presented as fold changes with respect to average values for control cells. Mean values (n = 3 ± SEM) are presented. Statistical significance tested by comparison to siCtrl#1. **(B)** Western blots showing the efficiency of CRISPR- Cas9-mediated depletion of TSG101 (using single gRNA) and its effect on the levels of VPS28 protein, as compared to control conditions (non-targeting gRNAs, gCtrl#1 or #2), in HepG2 cells. Vinculin used as a gel loading control. **(C)** qPCR results showing the expression of genes encoding indicated oxidative metabolism enzymes in HepG2 cells with CRISPR-Cas9-mediated depletion of TSG101, as compared to control cells presented as fold changes with respect to average values for control cells. Mean values (n = 4 ± SEM) are presented. Statistical significance tested by comparison to gCtrl#2. *P < 0.05, **P < 0.01, ***P < 0.001.

**Figure S2.**
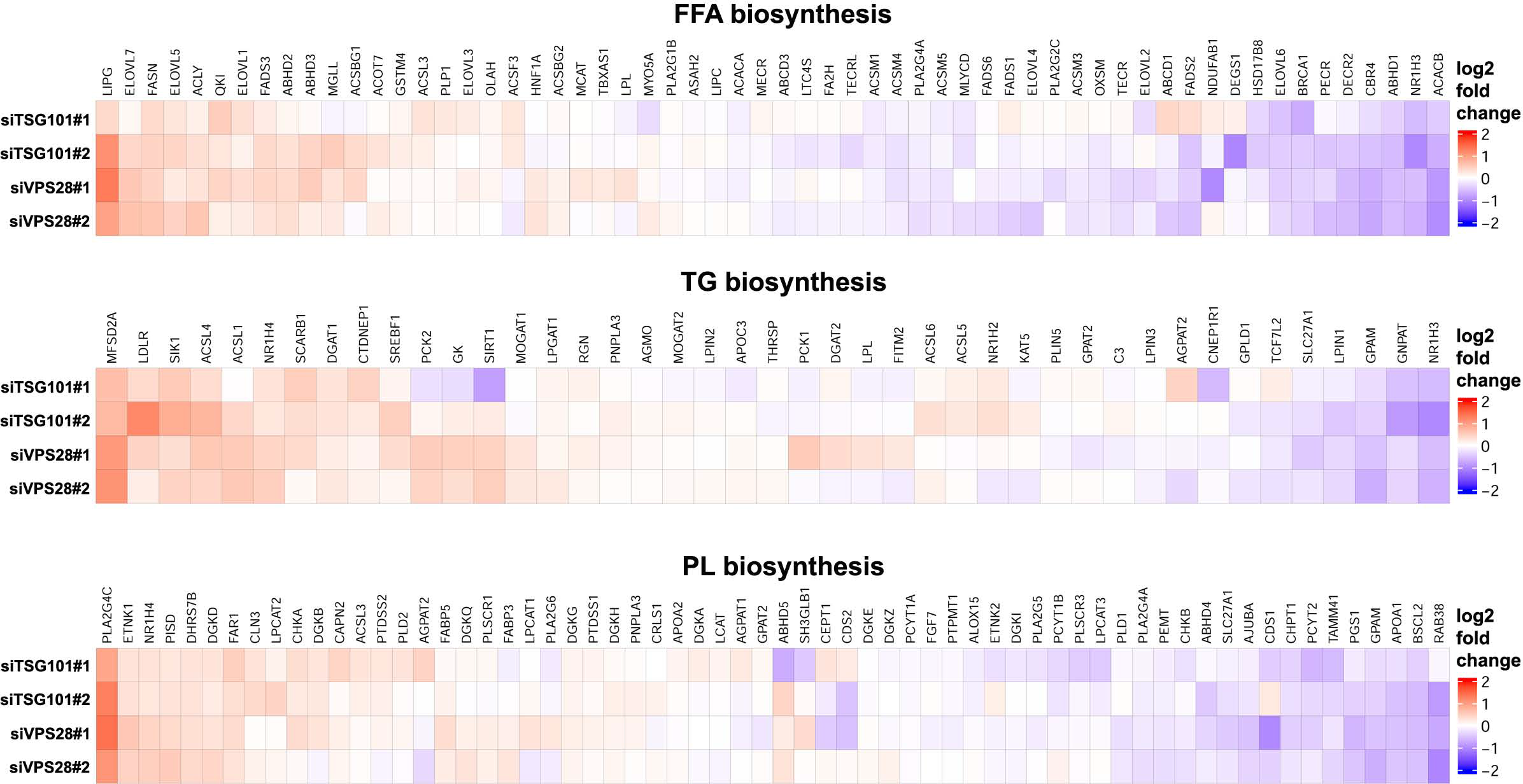
**Expression of several genes encoding enzymes involved in biosynthesis of fatty acid- containing lipids is increased in cells lacking ESCRT-I.** Heatmaps visualizing microarray results regarding the expression of genes encoding enzymes of fatty acid (FA), triglyceride (TG) or phospholipid (PL) biosynthesis in HEK293 cells after removal of ESCRT-I using two siRNAs for each component (siTSG101#1 or siTSG101#2, siVPS28#1 or siVPS28#2), as compared to control cells (treated with non-targeting siRNAs, Ctrl#1 or #2). Microarray data analysis was performed based on three independent experiments at three days post transfection with siRNAs (3 dpt).

**Figure S3.**
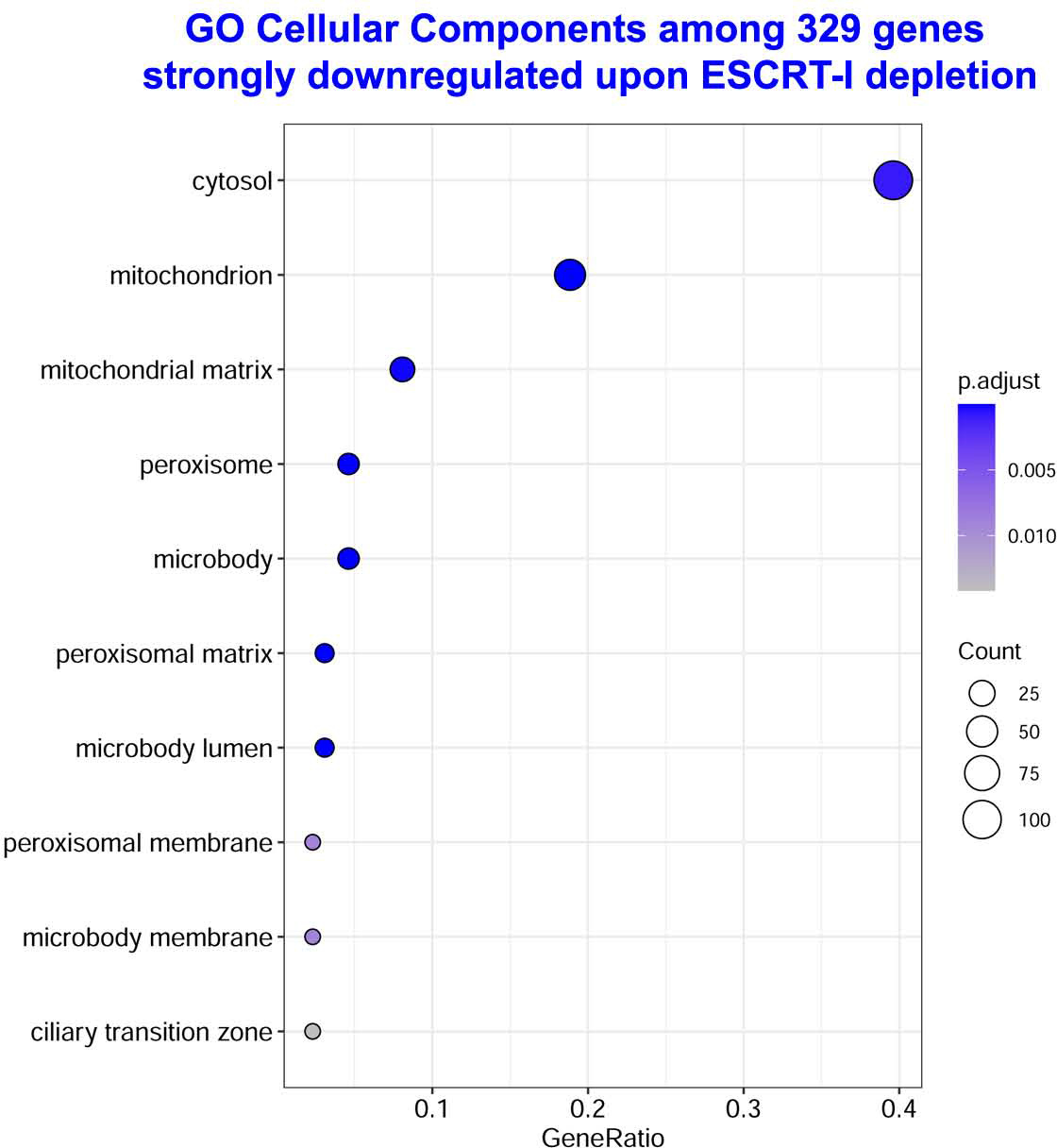
**Many genes with reduced expression upon ESCRT-I deficiency encode mitochondrial proteins.** Gene ontology (GO) analysis of top cellular compartments identified by annotation of genes detected in microarray experiments as those with strongly downregulated expression (log2 fold change ≤ -0.6; FDR < 0.05) in HEK293 cells after removal of ESCRT-I using two siRNAs for each component (siTSG101#1 or siTSG101#2, siVPS28#1 or siVPS28#2), as compared to control cells (treated with non-targeting siRNAs, Ctrl#1 or #2). Microarray data analysis was performed based on three independent experiments at three days post transfection with siRNAs (3 dpt).

**Figure S4.**
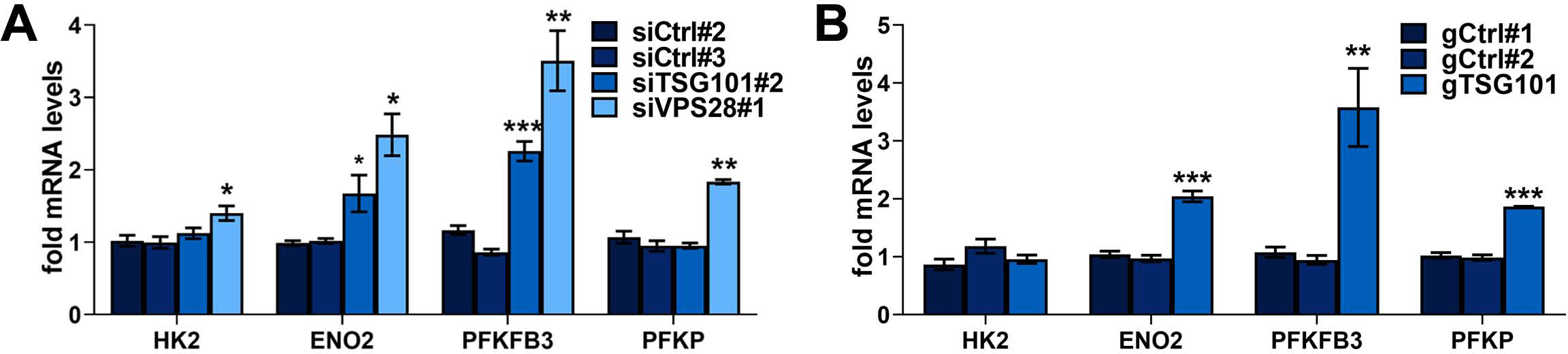
ESCRT-I dysfunction leads to increased expression of genes involved in glycolytic metabolism in HepG2 cells. (A-B) qPCR results showing the expression of genes encoding the indicated glycolytic metabolism enzymes in HepG2 cells with siRNA-mediated removal of ESCRT-I components using single siRNA for each component (siTSG101#2, siVPS28#1 in A) or in cells with CRISPR-Cas9-mediated depletion of TSG101 using single gRNA (gTsg101 in B), as compared to respective control cells (treated with non-targeting siRNAs, Ctrl#2 or #3 or gRNAs, gCtrl#1 or #2). The results presented as fold changes with respect to average values for control cells. Mean values (n = 3 ± SEM). Statistical significance tested by comparison to siCtrl#2 (in A) or gCtrl#1 (in B). *P < 0.05, **P < 0.01, ***P < 0.001.

**Figure S5.**
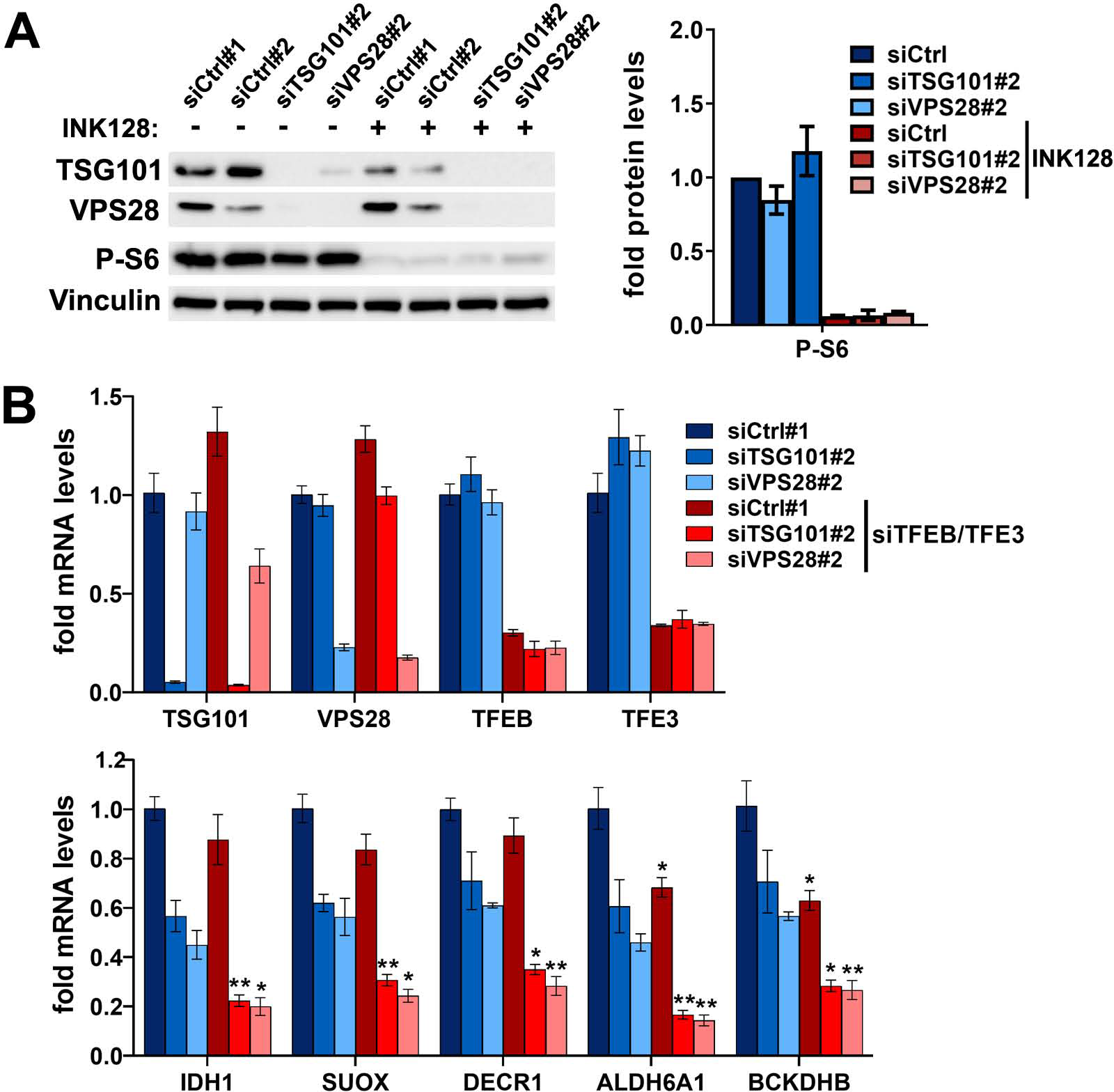
mTORC1 signaling does not contribute to reduced expression of genes encoding enzymes of amino acid or fatty acid oxidation upon ESCRT-I deficiency. (A) Representative western blots showing the levels of TSG101 and VPS28 proteins as well as phosphorylated S6 protein (P-S6) in control cells (siCtrl#1 or siCtrl#2) or cells depleted of ESCRT-I components (siTSG101#2 or siVPS28#2), treated with DMSO or 100 nM mTOR kinase inhibitor (INK128) for 48 h. The graph (right panel) shows protein levels as fold change with respect to averaged values measured for siCtrl#1 and #2 (siCtrl) assessed by densitometry analysis of western blotting bands. The analysis was performed at 3 dpt with vinculin used as a gel loading control. **(B)** qPCR results showing the expression of genes encoding components of ESCRT-I, TFEB/TFE3 transcription factors or the indicated enzymes of amino acid or fatty acid oxidation in control (siCtrl#1) or ESCRT-I-deficient (siTSG101#2 or siVPS28#2) HEK293 cells with single depletion or with co-depletion of TFEB and TFE3 (siTFEB/TFE3). The results presented as fold changes with respect to siCtrl#1 at 4 dpt. Mean values (n = 4 ± SEM) are presented. Statistical significance tested by comparison of cells lacking only TSG101 or VPS28 to those lacking these proteins and TFEB/TFE3. *P < 0.05, **P < 0.01.

**Figure S6.**
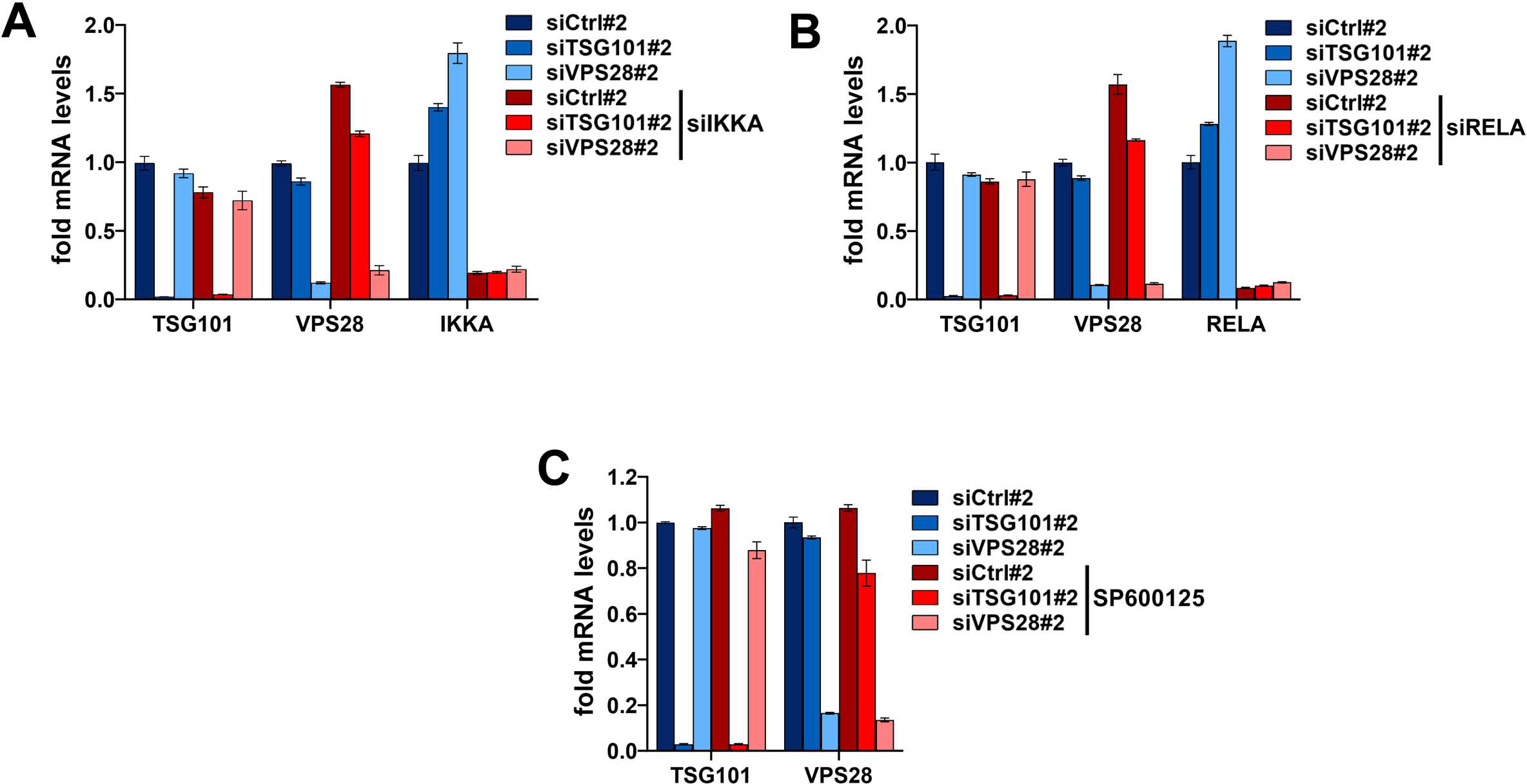
Simultaneous depletion of IKKA or RELA or treatment with SP600125 do not interfere with efficient depletion of ESCRT-I components. (A-B) qPCR results showing the efficiencies of silencing the expression of genes encoding the indicated proteins in cells treated with siRNAs targeting the ESCRT-I components (siTSG101#2 or siVPS28#2) or regulators of NFκB signaling (siIKKA or siRELA) as compared to control cells (siCtrl#2) at four days post siRNA transfection (4 dpt). Mean values (n = 4 ± SEM in A or n = 3 ± SEM in B) are presented. (C) qPCR results showing the efficiencies of silencing the expression of genes encoding TSG101 or VPS28 proteins in cells treated with siTSG101#2 or siVPS28#2 as compared to controls cells (siCtrl#2) upon 72 h treatment with DMSO or 50 μM SP600125 compound.

## REFERENCES

1. Vander Heiden MG, Cantley LC, Thompson CB (2009) Understanding the Warburg effect: the metabolic requirements of cell proliferation. Science, 324:1029–1033.

2. Ye Z, Wang S, Zhang C, Zhao Y (2020) Coordinated Modulation of Energy Metabolism and Inflammation by Branched-Chain Amino Acids and Fatty Acids. Front Endocrinol (Lausanne), 11:617.

3. Houten SM, Violante S, Ventura FV, Wanders RJ (2016) The Biochemistry and Physiology of Mitochondrial Fatty Acid beta-Oxidation and Its Genetic Disorders. Annu Rev Physiol, 78:23–44.

4. Tanosaki S, Tohyama S, Kishino Y, Fujita J, Fukuda K (2021) Metabolism of human pluripotent stem cells and differentiated cells for regenerative therapy: a focus on cardiomyocytes. Inflamm Regen, 41:5.

5. Riester M, Xu Q, Moreira A, Zheng J, Michor F, Downey RJ (2018) The Warburg effect: persistence of stem-cell metabolism in cancers as a failure of differentiation. Ann Oncol, 29:264–270.

6. Kornberg MD (2020) The immunologic Warburg effect: Evidence and therapeutic opportunities in autoimmunity. Wiley Interdiscip Rev Syst Biol Med, 12:e1486.

7. Krishnan J, Suter M, Windak R, Krebs T, Felley A, Montessuit C, Tokarska-Schlattner M, Aasum E, Bogdanova A, Perriard E, Perriard JC, Larsen T, Pedrazzini T, Krek W (2009) Activation of a HIF1alpha-PPARgamma axis underlies the integration of glycolytic and lipid anabolic pathways in pathologic cardiac hypertrophy. Cell Metab, 9:512–524.

8. Li Z, Zhang H (2016) Reprogramming of glucose, fatty acid and amino acid metabolism for cancer progression. Cell Mol Life Sci, 73:377–392.

9. Lawrence RE, Zoncu R (2019) The lysosome as a cellular centre for signalling, metabolism and quality control. Nat Cell Biol, 21:133–142.

10. Deus CM, Yambire KF, Oliveira PJ, Raimundo N (2020) Mitochondria-Lysosome Crosstalk: From Physiology to Neurodegeneration. Trends Mol Med, 26:71–88.

11. Wrobel M, Cendrowski J, Szymanska E, Grebowicz-Maciukiewicz M, Budick-Harmelin N, Macias M, Szybinska A, Mazur M, Kolmus K, Goryca K, Dabrowska M, Paziewska A, Mikula M, Miaczynska M (2022) ESCRT-I fuels lysosomal degradation to restrict TFEB/TFE3 signaling via the Rag- mTORC1 pathway. Life Sci Alliance, 5.

12. von Zastrow M, Sorkin A (2021) Mechanisms for Regulating and Organizing Receptor Signaling by Endocytosis. Annu Rev Biochem, 90:709–737.

13. Gilleron J, Zeigerer A (2023) Endosomal trafficking in metabolic homeostasis and diseases. Nat Rev Endocrinol, 19:28–45.

14. Adriaenssens E, Ferrari L, Martens S (2022) Orchestration of selective autophagy by cargo receptors. Curr Biol, 32:R1357–R1371.

15. Palm W, Thompson CB (2017) Nutrient acquisition strategies of mammalian cells. Nature, 546:234–242.

16. Shin HR, Zoncu R (2020) The Lysosome at the Intersection of Cellular Growth and Destruction. Dev Cell, 54:226–238.

17. Castellano BM, Thelen AM, Moldavski O, Feltes M, van der Welle RE, Mydock-McGrane L, Jiang X, van Eijkeren RJ, Davis OB, Louie SM, Perera RM, Covey DF, Nomura DK, Ory DS, Zoncu R (2017) Lysosomal cholesterol activates mTORC1 via an SLC38A9-Niemann-Pick C1 signaling complex. Science, 355:1306–1311.

18. Zoncu R, Bar-Peled L, Efeyan A, Wang S, Sancak Y, Sabatini DM (2011) mTORC1 senses lysosomal amino acids through an inside-out mechanism that requires the vacuolar H(+)-ATPase. Science, 334:678–683.

19. Vietri M, Radulovic M, Stenmark H (2020) The many functions of ESCRTs. Nat Rev Mol Cell Biol, 21:25-42.

20. Olmos Y (2022) The ESCRT Machinery: Remodeling, Repairing, and Sealing Membranes. Membranes (Basel), 12.

21. Szymanska E, Budick-Harmelin N, Miaczynska M (2018) Endosomal "sort" of signaling control: The role of ESCRT machinery in regulation of receptor-mediated signaling pathways. Semin Cell Dev Biol, 74:11–20.

22. Filimonenko M, Stuffers S, Raiborg C, Yamamoto A, Malerod L, Fisher EM, Isaacs A, Brech A, Stenmark H, Simonsen A (2007) Functional multivesicular bodies are required for autophagic clearance of protein aggregates associated with neurodegenerative disease. J Cell Biol, 179:485–500.

23. Zhen Y, Spangenberg H, Munson MJ, Brech A, Schink KO, Tan KW, Sorensen V, Wenzel EM, Radulovic M, Engedal N, Simonsen A, Raiborg C, Stenmark H (2020) ESCRT-mediated phagophore sealing during mitophagy. Autophagy, 16:826–841.

24. Hammerling BC, Najor RH, Cortez MQ, Shires SE, Leon LJ, Gonzalez ER, Boassa D, Phan S, Thor A, Jimenez RE, Li H, Kitsis RN, Dorn GW, II, Sadoshima J, Ellisman MH, Gustafsson AB (2017) A Rab5 endosomal pathway mediates Parkin-dependent mitochondrial clearance. Nat Commun, 8:14050.

25. Wegner CS, Rodahl LM, Stenmark H (2011) ESCRT proteins and cell signalling. Traffic, 12:1291–1297.

26. Banach-Orlowska M, Jastrzebski K, Cendrowski J, Maksymowicz M, Wojciechowska K, Korostynski M, Moreau D, Gruenberg J, Miaczynska M (2018) The topology of the lymphotoxin beta receptor that accumulates upon endolysosomal dysfunction dictates the NF-kappaB signaling outcome. J Cell Sci, 131:jcs218883.

27. Maminska A, Bartosik A, Banach-Orlowska M, Pilecka I, Jastrzebski K, Zdzalik-Bielecka D, Castanon I, Poulain M, Neyen C, Wolinska-Niziol L, Torun A, Szymanska E, Kowalczyk A, Piwocka K, Simonsen A, Stenmark H, Furthauer M, Gonzalez-Gaitan M, Miaczynska M (2016) ESCRT proteins restrict constitutive NF-kappaB signaling by trafficking cytokine receptors. Sci Signal, 9:ra8.

28. Kolmus K, Erdenebat P, Szymanska E, Stewig B, Goryca K, Derezinska-Wolek E, Szumera-Cieckiewicz A, Brewinska-Olchowik M, Piwocka K, Prochorec-Sobieszek M, Mikula M, Miaczynska M (2021) Concurrent depletion of Vps37 proteins evokes ESCRT-I destabilization and profound cellular stress responses. J Cell Sci, 134:jcs250951.

29. Woodfield SE, Graves HK, Hernandez JA, Bergmann A (2013) De-regulation of JNK and JAK/STAT signaling in ESCRT-II mutant tissues cooperatively contributes to neoplastic tumorigenesis. PLoS One, 8:e56021.

30. Oshima R, Hasegawa T, Tamai K, Sugeno N, Yoshida S, Kobayashi J, Kikuchi A, Baba T, Futatsugi A, Sato I, Satoh K, Takeda A, Aoki M, Tanaka N (2016) ESCRT-0 dysfunction compromises autophagic degradation of protein aggregates and facilitates ER stress-mediated neurodegeneration via apoptotic and necroptotic pathways. Sci Rep, 6:24997.

31. Planavila A, Laguna JC, Vazquez-Carrera M (2005) Nuclear factor-kappaB activation leads to down- regulation of fatty acid oxidation during cardiac hypertrophy. J Biol Chem, 280:17464–17471.

32. Londhe P, Yu PY, Ijiri Y, Ladner KJ, Fenger JM, London C, Houghton PJ, Guttridge DC (2018) Classical NF-kappaB Metabolically Reprograms Sarcoma Cells Through Regulation of Hexokinase 2. Front Oncol, 8:104.

33. Papa S, Choy PM, Bubici C (2019) The ERK and JNK pathways in the regulation of metabolic reprogramming. Oncogene, 38:2223–2240.

34. Sekar R, et al (2022) Vps37a regulates hepatic glucose production by controlling glucagon receptor localization to endosomes. Cell Metab, 34:1824–1842 e1829.

35. Settembre C, De Cegli R, Mansueto G, Saha PK, Vetrini F, Visvikis O, Huynh T, Carissimo A, Palmer D, Klisch TJ, Wollenberg AC, Di Bernardo D, Chan L, Irazoqui JE, Ballabio A (2013) TFEB controls cellular lipid metabolism through a starvation-induced autoregulatory loop. Nat Cell Biol, 15:647–658.

36. Mansueto G, et al (2017) Transcription Factor EB Controls Metabolic Flexibility during Exercise. Cell Metab, 25:182–196.

37. Bache KG, Slagsvold T, Cabezas A, Rosendal KR, Raiborg C, Stenmark H (2004) The growth-regulatory protein HCRP1/hVps37A is a subunit of mammalian ESCRT-I and mediates receptor down-regulation. Mol Biol Cell, 15:4337–4346.

38. Geisbrecht BV, Gould SJ (1999) The human PICD gene encodes a cytoplasmic and peroxisomal NADP(+)-dependent isocitrate dehydrogenase. J Biol Chem, 274:30527–30533.

39. Nassar ZD, Mah CY, Dehairs J, Burvenich IJ, Irani S, Centenera MM, Helm M, Shrestha RK, Moldovan M, Don AS, Holst J, Scott AM, Horvath LG, Lynn DJ, Selth LA, Hoy AJ, Swinnen JV, Butler LM (2020) Human DECR1 is an androgen-repressed survival factor that regulates PUFA oxidation to protect prostate tumor cells from ferroptosis. Elife, 9.

40. Reschke S, Niks D, Wilson H, Sigfridsson KG, Haumann M, Rajagopalan KV, Hille R, Leimkuhler S (2013) Effect of exchange of the cysteine molybdenum ligand with selenocysteine on the structure and function of the active site in human sulfite oxidase. Biochemistry, 52:8295–8303.

41. Quental S, Macedo-Ribeiro S, Matos R, Vilarinho L, Martins E, Teles EL, Rodrigues E, Diogo L, Garcia P, Eusebio F, Gaspar A, Sequeira S, Furtado F, Lanca I, Amorim A, Prata MJ (2008) Molecular and structural analyses of maple syrup urine disease and identification of a founder mutation in a Portuguese Gypsy community. Mol Genet Metab, 94:148–156.

42. Tanner JJ (2015) SAXS fingerprints of aldehyde dehydrogenase oligomers. Data Brief, 5:745–751.

43. Ravnskjaer K, Frigerio F, Boergesen M, Nielsen T, Maechler P, Mandrup S (2010) PPARdelta is a fatty acid sensor that enhances mitochondrial oxidation in insulin-secreting cells and protects against fatty acid-induced dysfunction. J Lipid Res, 51:1370–1379.

44. Fowler SD, Greenspan P (1985) Application of Nile red, a fluorescent hydrophobic probe, for the detection of neutral lipid deposits in tissue sections: comparison with oil red O. J Histochem Cytochem, 33:833–836.

45. Qiu B, Simon MC (2016) BODIPY 493/503 Staining of Neutral Lipid Droplets for Microscopy and Quantification by Flow Cytometry. Bio Protoc, 6.

46. Diaz G, Melis M, Batetta B, Angius F, Falchi AM (2008) Hydrophobic characterization of intracellular lipids in situ by Nile Red red/yellow emission ratio. Micron, 39:819–824.

47. Wang G, Yu Y, Wang YZ, Zhu ZM, Yin PH, Xu K (2020) Effects and mechanisms of fatty acid metabolism-mediated glycolysis regulated by betulinic acid-loaded nanoliposomes in colorectal cancer. Oncol Rep, 44:2595–2609.

48. Israel A (2010) The IKK complex, a central regulator of NF-kappaB activation. Cold Spring Harb Perspect Biol, 2:a000158.

49. Kauppinen A, Suuronen T, Ojala J, Kaarniranta K, Salminen A (2013) Antagonistic crosstalk between NF-kappaB and SIRT1 in the regulation of inflammation and metabolic disorders. Cell Signal, 25:1939–1948.

50. Kunsch C, Rosen CA (1993) NF-kappa B subunit-specific regulation of the interleukin-8 promoter. Mol Cell Biol, 13:6137–6146.

51. Kim BG, Sung JS, Jang Y, Cha YJ, Kang S, Han HH, Lee JH, Cho NH (2019) Compression-induced expression of glycolysis genes in CAFs correlates with EMT and angiogenesis gene expression in breast cancer. Commun Biol, 2:313.

52. Umebashi K, Tokito A, Yamamoto M, Jougasaki M (2018) Interleukin-33 induces interleukin-8 expression via JNK/c-Jun/AP-1 pathway in human umbilical vein endothelial cells. PLoS One, 13:e0191659.

53. Rasheed Z, Akhtar N, Haqqi TM (2011) Advanced glycation end products induce the expression of interleukin-6 and interleukin-8 by receptor for advanced glycation end product-mediated activation of mitogen-activated protein kinases and nuclear factor-kappaB in human osteoarthritis chondrocytes. Rheumatology (Oxford), 50:838–851.

54. Jo MS, Yang HW, Park JH, Shin JM, Park IH (2023) Glycolytic reprogramming is involved in tissue remodeling on chronic rhinosinusitis. PLoS One, 18:e0281640.

55. Mehla K, Singh PK (2019) Metabolic Regulation of Macrophage Polarization in Cancer. Trends Cancer, 5:822–834.

56. Nishimura K, Fukuda A, Hisatake K (2019) Mechanisms of the Metabolic Shift during Somatic Cell Reprogramming. Int J Mol Sci, 20.

57. Antonescu CN, McGraw TE, Klip A (2014) Reciprocal regulation of endocytosis and metabolism. Cold Spring Harb Perspect Biol, 6:a016964.

58. Gilleron J, Gerdes JM, Zeigerer A (2019) Metabolic regulation through the endosomal system. Traffic, 20:552-570.

59. Jin L, Huo Y, Zheng Z, Jiang X, Deng H, Chen Y, Lian Q, Ge R, Deng H (2014) Down-regulation of Ras-related protein Rab 5C-dependent endocytosis and glycolysis in cisplatin-resistant ovarian cancer cell lines. Mol Cell Proteomics, 13:3138–3151.

60. Willinger T, Staron M, Ferguson SM, De Camilli P, Flavell RA (2015) Dynamin 2-dependent endocytosis sustains T-cell receptor signaling and drives metabolic reprogramming in T lymphocytes. Proc Natl Acad Sci U S A, 112:4423–4428.

61. Zeigerer A, et al (2015) Regulation of liver metabolism by the endosomal GTPase Rab5. Cell Rep, 11:884-892.

62. Brass EP (1994) Overview of coenzyme A metabolism and its role in cellular toxicity. Chem Biol Interact, 90:203–214.

63. Vanweert F, Schrauwen P, Phielix E (2022) Role of branched-chain amino acid metabolism in the pathogenesis of obesity and type 2 diabetes-related metabolic disturbances BCAA metabolism in type 2 diabetes. Nutr Diabetes, 12:35.

64. Lu L, Chen Y, Zhu Y (2017) The molecular basis of targeting PFKFB3 as a therapeutic strategy against cancer. Oncotarget, 8:62793–62802.

65. Hu ZY, Xiao L, Bode AM, Dong Z, Cao Y (2014) Glycolytic genes in cancer cells are more than glucose metabolic regulators. J Mol Med (Berl), 92:837–845.

66. Loi M, Raimondi A, Morone D, Molinari M (2019) ESCRT-III-driven piecemeal micro-ER-phagy remodels the ER during recovery from ER stress. Nat Commun, 10:5058.

67. Wang J, Fang N, Xiong J, Du Y, Cao Y, Ji WK (2021) An ESCRT-dependent step in fatty acid transfer from lipid droplets to mitochondria through VPS13D-TSG101 interactions. Nat Commun, 12:1252.

68. Perera RM, Di Malta C, Ballabio A (2019) MiT/TFE Family of Transcription Factors, Lysosomes, and Cancer. Annu Rev Cancer Biol, 3:203–222.

69. Remels AH, Gosker HR, Verhees KJ, Langen RC, Schols AM (2015) TNF-alpha-induced NF-kappaB activation stimulates skeletal muscle glycolytic metabolism through activation of HIF-1alpha. Endocrinology, 156:1770–1781.

70. Han D, Wei W, Chen X, Zhang Y, Wang Y, Zhang J, Wang X, Yu T, Hu Q, Liu N, You Y (2015) NF-kappaB/RelA-PKM2 mediates inhibition of glycolysis by fenofibrate in glioblastoma cells. Oncotarget, 6:26119–26128.

71. Mauro C, Leow SC, Anso E, Rocha S, Thotakura AK, Tornatore L, Moretti M, De Smaele E, Beg AA, Tergaonkar V, Chandel NS, Franzoso G (2011) NF-kappaB controls energy homeostasis and metabolic adaptation by upregulating mitochondrial respiration. Nat Cell Biol, 13:1272–1279.

72. Zhdanov AV, Dmitriev RI, Papkovsky DB (2011) Bafilomycin A1 activates respiration of neuronal cells via uncoupling associated with flickering depolarization of mitochondria. Cell Mol Life Sci, 68:903–917.

73. Shum LC, White NS, Mills BN, Bentley KL, Eliseev RA (2016) Energy Metabolism in Mesenchymal Stem Cells During Osteogenic Differentiation. Stem Cells Dev, 25:114–122.

74. Zheng X, Boyer L, Jin M, Mertens J, Kim Y, Ma L, Ma L, Hamm M, Gage FH, Hunter T (2016) Metabolic reprogramming during neuronal differentiation from aerobic glycolysis to neuronal oxidative phosphorylation. Elife, 5.

75. Sandoval IT, Delacruz RG, Miller BN, Hill S, Olson KA, Gabriel AE, Boyd K, Satterfield C, Van Remmen H, Rutter J, Jones DA (2017) A metabolic switch controls intestinal differentiation downstream of Adenomatous polyposis coli (APC). Elife, 6.

76. Doench JG, Fusi N, Sullender M, Hegde M, Vaimberg EW, Donovan KF, Smith I, Tothova Z, Wilen C, Orchard R, Virgin HW, Listgarten J, Root DE (2016) Optimized sgRNA design to maximize activity and minimize off-target effects of CRISPR-Cas9. Nat Biotechnol, 34:184–191.

77. Barde I, Salmon P, Trono D (2010) **Production and titration of lentiviral vectors.** *Curr Protoc Neurosci*, **Chapter** 4:Unit 4 21.

78. Pera J, Korostynski M, Golda S, Piechota M, Dzbek J, Krzyszkowski T, Dziedzic T, Moskala M, Przewlocki R, Szczudlik A, Slowik A (2013) Gene expression profiling of blood in ruptured intracranial aneurysms: in search of biomarkers. J Cereb Blood Flow Metab, 33:1025–1031.

79. Yu G, Wang LG, Han Y, He QY (2012) clusterProfiler: an R package for comparing biological themes among gene clusters. OMICS, 16:284–287.

80. Gu Z, Eils R, Schlesner M (2016) Complex heatmaps reveal patterns and correlations in multidimensional genomic data. Bioinformatics, 32:2847–2849.

81. Bligh EG, Dyer WJ (1959) A rapid method of total lipid extraction and purification. Can J Biochem Physiol, 37:911–917.

